# Identification of key proteins involved in stickleback environmental adaption with system-level analysis

**DOI:** 10.1101/2020.02.11.943522

**Authors:** Martina Hall, Dietmar Kültz, Eivind Almaas

## Abstract

Using abundance measurements of 1,490 proteins from four separate populations of three-spined sticklebacks, we implemented a system-level approach to correlate proteome dynamics with environmental salinity and temperature and the fish’s population and morphotype. We identified sets of robust and accurate fingerprints that predict environmental salinity, temperature, morphotype and the population sample origin, observing that proteins with specific functions are enriched in these fingerprints. Highly apparent functions represented in all fingerprints include ion transport, proteostasis, growth, and immunity, suggesting that these functions are most diversified in populations inhabiting different environments.

Applying a differential network approach, we analyzed the network of protein interactions that differs between populations. Looking at specific population combinations of differential interaction, we identify sets of connected proteins. We find that these sets and their corresponding enriched functions reflect key processes that have diverged between the four populations. Moreover, the extent of divergence, i.e. the number of enriched functions that differ between populations, is highest when all three environmental parameters are different between two populations. Key nodes in the differential interaction network signify functions that are also inherent in the fingerprints, most prominently proteostasis-related functions. However, the differential interaction network also reveals additional functions that have diverged between populations, notably cytoskeletal organization and morphogenesis.

Having such a large proteomic dataset, the strength of these analyses is that the results are purely data-driven, not based on previous findings and hypotheses about adaptation. With such an unbiased approach applied on a large proteomic dataset, we find the strongest signals given by the data, making it possible to develop more discriminatory and complex biomarkers for specific contexts of interest.

## Introduction

The three-spined stickleback is a particularly interesting specie when studying environmental adaption. They can inhabit very different environments; from cold marine water, to cold freshwater lakes, to warm brackish lagoons, and also display different morphotypes. Many studies have been undertaken with the intent of gaining a deep understanding of how this fish is able to adapt to such different environments, mostly focused on adaption to salinity and the morphotype of the fish. Nevertheless, large-scale studies that bridge the gap between the genome and organismal phenotypes by comparative analyses of molecular phenotypes (the proteome) in divergently adapted populations are still in their infancy. Using proteomics, one can study adaptation through the molecular building blocks, representing the link between genetic information (specific genomic loci) and phenotypic variation at higher levels of organization, including organismal structure and function.

Here, we present a system-level data-driven analysis to shed light on this question using a recent large-scale proteomics dataset from three-spine stickleback gills^1^. The authors conducted measurements on a total of 96 fish, consisting of groups of 24 fish sampled from four separate locations characterized by the following environmental parameters: cold, brackish water (BW) in Westchester Lagoon, Alaska (AK), cold salt water (SW) in Bodega Harbor, California (BH), cold freshwater (FW) in Lake Solano, California (LS) and warm brackish water in Laguna de la Bocana del Rosario, Mexico (RE). The sampled fish also represent different morphotypes, where fish from the AK and BH populations are of the trachurus morphotype (large body size, fully plated) and fish from LS and RE populations are of the leiurus morphotype (small, low-plated). This dataset enables more stringent statistical analyses and is more suitable for network-biology approaches because it relies on identical measurement parameters (transitions, precursors, and proteins) that are quantified for every sample. In contrast, conventional proteomics methods such as shotgun LCMS sample data stochastically and result in heterogeneous data structures. On the other hand, targeted proteomics approaches such as selective, parallel, and multiple reaction monitoring are not suitable for systems-scale analyses because they only monitor a limited number of proteins.

To understand how these fish have adapted to such different environments, we explore the data for patterns of protein abundance specific for the environmental traits. Using the protein abundance measured for each fish, we first apply the Least Absolute Shrinkage and Selection Operator (LASSO) method^2^, an unsupervised statistical learning method for variable selection and prediction in high dimensions, to find robust protein fingerprints for the environmental traits. With this method, we identify small sets of proteins that are the most important for separation between the traits, where the specific combination of these proteins give rise to accurate statistical prediction models for the environmental traits.

In addition to identifying fingerprint proteins specific for the environmental traits, we use a network approach that is based on comparing pairs of proteins across the different stickleback habitats. Analyzing the properties of the resulting network, we get a system-level overview of how the proteins work together and affect each other across the habitats. In previous work^3^, s-core network-peeling was performed on parts of these data, where they found differences in the network structure of protein interactions for fresh and marine water. Here, we extend these analyses by investigating differential interactions for the four populations. Applying the Co-expression Differential Network Analysis approach^4^ (CoDiNA), we first create correlative protein interaction networks for each population, where the proteins are represented as nodes and the correlation between proteins are represented as links. The four networks are subsequently merged into one using the CoDiNA framework, where we extract links with opposite correlation structure for at least two populations. With this method, we identify differential protein interactions between the populations and key proteins possibly affecting the system-level adaptation to different environments.

As massive amounts of data are generated by omics approaches and improvement of omics technologies, it necessitates parallel development and use of computational analysis approaches for rapidly identifying the biologically most meaningful features embedded in those data, extracting the most relevant information, and simplifying visual illustration of the main conclusions supported by large and complex datasets. One of the needs for interpreting omics-type datasets that are comprehensive and include large differences between groups is to condense these differences down to the smallest set that still retains most of the pertinent biological information/ interpretation. This condensation is usually investigator-driven, manual and, therefore, biased. Taking on a mathematical / statistical procedure, our approach provides an unbiased alternative purely based on the data. We make no hypothesis about the proteins, but simply extract the strongest signals present. Using existing methods for high-dimensional data analysis, we analyze the large proteomics dataset with unsupervised learning methods, and the results are hence data-driven.

## Results

With the publication of the protein-abundance data^1^, the authors identified proteins with elevated and reduced abundance levels using univariate statistics and separately testing each protein for differences in abundance in one population against the others. This basic set of tests identified proteins that were present with significantly elevated or reduced abundance in a specific population compared to the other populations, and sets of these proteins that were enriched for biological functions specific for a fish environment.

In this paper we further and significantly expand these analyses by presenting results from a system-level approach, not only looking at single-protein differences among the populations, but the differences in sets of interactions between proteins and in the combination of proteins that statistically explain the ability of the fish to inhabit these different environments. We first present results from a Principal Component Analysis (PCA), where we investigate variability among samples from the four populations. We continue by using a statistical variable-selection method to identify sets of proteins, where the combination of these proteins give rise to accurate prediction models for these environments. Finally, we employ a network-based approach, identifying sets of interacting proteins that are specific and differential between the population habitats.

### Dimension-reduction analysis with PCA

In order to generate a system-level understanding of the sets of differentially expressed proteins involved in the adaptive changes in the four populations, we employ the much used Principal Component Analysis (PCA)^5^ as an initial approach. Using PCA, we identify principal components (PCs) that are linear combinations of proteins according to their contribution to the variance in the data. Each protein will contribute with a loading score to each PC, where the first PC captures the most variability among the data, with subsequent PCs explaining decreasing amounts of variability. If the number of PCs necessary to capture a substantial amount of variation between the different populations is small, this will provide a robust method for visualizing the data and also identify sets of proteins central in the adaptation processes for the four environments.

Figure 1 shows the results using the first three principal components. The first thing to observe is that it is necessary to use a large number of principal components to explain the inherent variation in the data (Figure 1 D). The cumulative scree plot shows that it is necessary to use 25 of the PCs to explain more than 75% of the variability in the data. On the other hand, we find that PC2 is able to provide a clear separation between the RE population and the three others. Additionally, there seems to be some degree of clustering of the populations when viewing them in PC1 and PC3 dimension. However, the loading vectors (Figure 1, panels A)-C), grey arrows) with the largest contributions for each PC have scores only in [−0.06, 0.06]. This indicates that these proteins explain only a small fraction of the PCs. Consequently, it is not possible to draw reliable conclusions about the proteins participating in the PCs to explain the observed groupings, and we need other methods to determine which proteins are involved in the stickleback adaptation to the four different environments.

**Figure 1.**
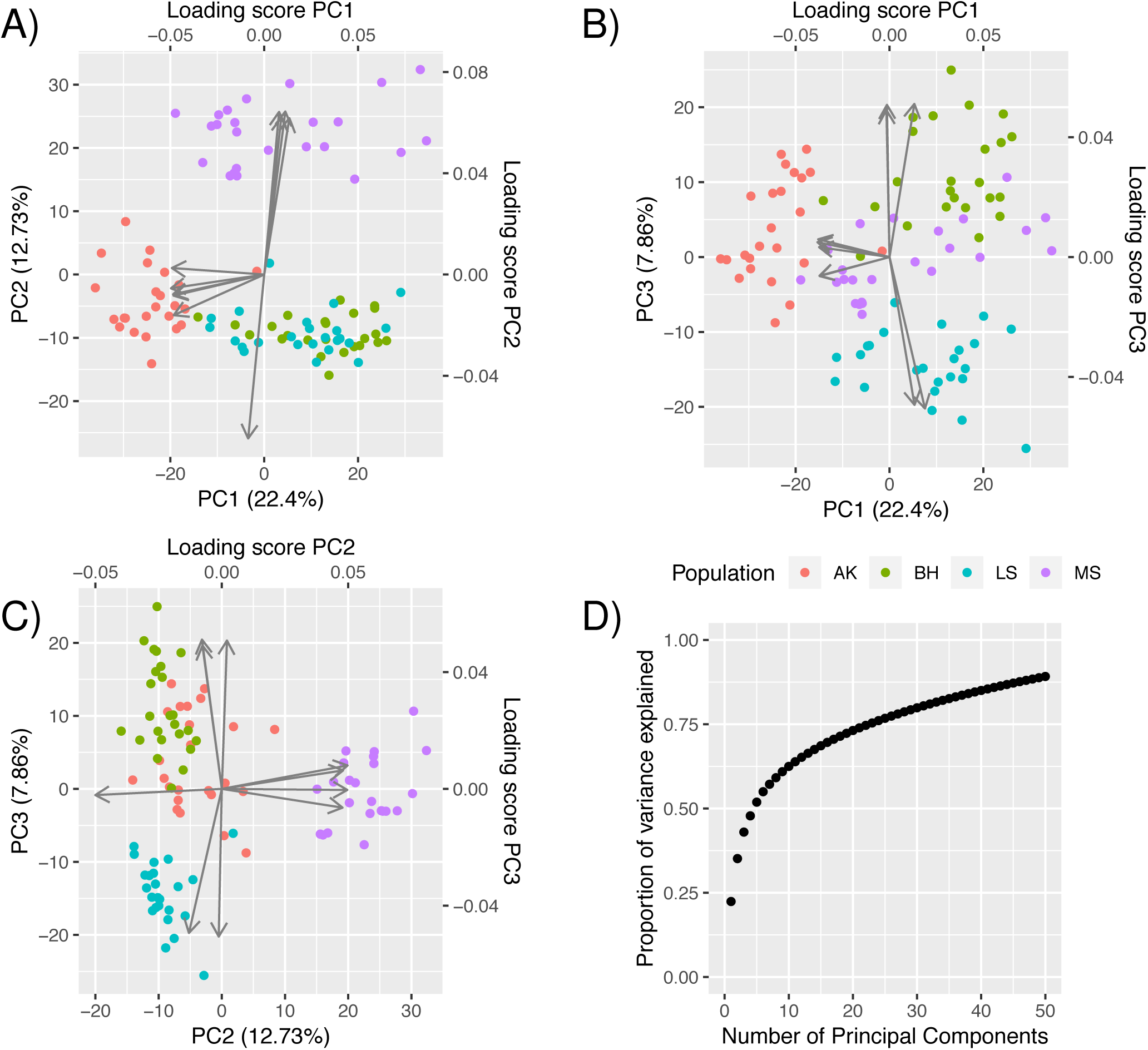
PCA to study protein expression variation among the four stickleback populations. Sample points are colored according their population and plotted in **A)** PC1 vs. PC2, **B)** PC1 vs. PC3, and **C)** PC2 vs. PC3. The top-5 loading vectors for each PC are plotted according to the shown principal components, with axis scales marked on the top and right side of each panel. **D)** The cumulative proportion of variance in the protein abundance data explained by the number of PCs included.

### Identification of fingerprint proteins for the four populations using LASSO

Taking a different approach for dimension reduction, we apply the LASSO^2^ machine learning method that performs prediction and dimensionality reduction via subset selection. Using the LASSO method, we identify a set of “fingerprint” proteins, where the combination of this specific set of proteins give rise to an accurate statistical prediction model for the fish population (see Methods section for details). In this way, the LASSO approach differs from standard univariate analysis where, instead of testing each protein, one considers all proteins at the same time and a resulting model consists of a specific combination of proteins that performs the best at explaining the differences between the populations.

Using LASSO with a multinomial regression model^6^ and taking all of the 1,490 proteins as covariates with the fish population (AK, BH, LS, or RE) as responses, our regression model only needs 13 of the proteins (0.87%) to predict which population each sample belongs to with 100% accuracy (see Tab. 1 for coefficient estimates). This set of proteins can hence be viewed as a “fingerprint” for the populations, where the combination of these proteins is specific for population determination. Thus, conducting new sample measurements of these 13 proteins and running them through the model, this specific linear combination of the 13 proteins is able to identify the population of the new fish sample.

Taking a closer look at the measured abundance levels of the fingerprint proteins separated on the four populations, Fig 2 demonstrates that each of the selected proteins is specific to only a single population in its abundance distribution. The four proteins with orange color (AK population) have higher abundance levels for population AK than for the others, thus contributing strongly in separating between AK and the other populations. Similarly for population BH (blue) and LS (green), where high protein abundance levels of the proteins will contribute strongly in their population separations. For population RE (pink), the pattern is different in that it consists of both high (G3NT01 and G3PSB5) and low (G3NZU2) abundance levels for the separation of RE from the other populations. Thus, the regression model represented by the proteins in Fig 2 has a direct visual interpretation. In the case that proteomics data from a new fish are provided that contain measurements of the 13 fingerprint proteins, the following situation would occur if the fish originated from the AK population: 1) Values close to the orange boxes would be indicative of population AK, 2) values lower than the blue and 3) green boxes for these proteins are evidence against an origin being in the BH and LS populations. Finally, 4) values lower than the pink boxes for G3NT01 and G3PSB5 and higher than the box for G3NZU2 is evidence against the RE population. Hence, for this gedankenexperiment the fingerprint profile presents strong evidence that the sampled fish originated from the AK population.

**Figure 2.**
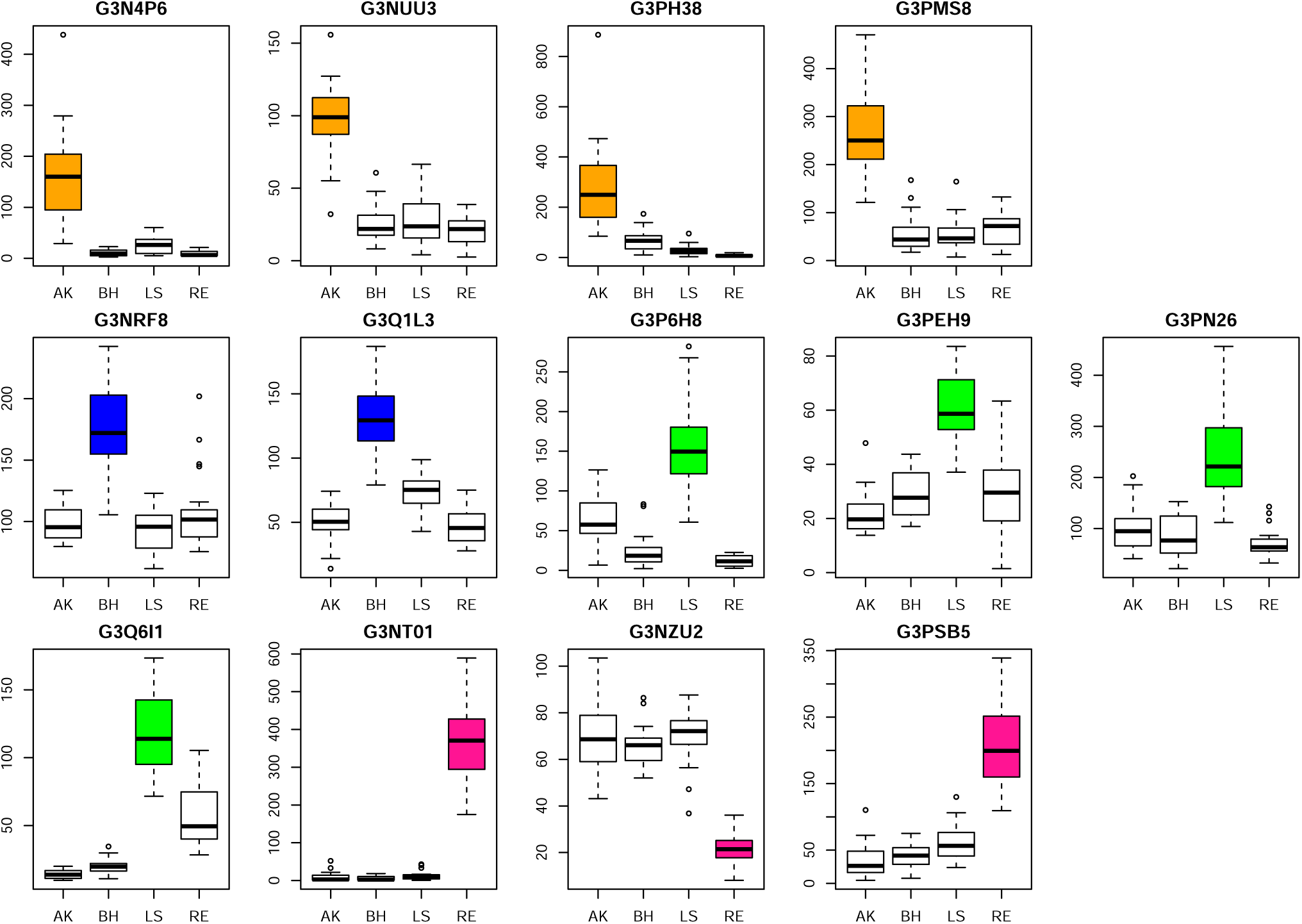
Abundance plot of the population fingerprint proteins. Each box contains the quartiles and the median of the measured abundance levels for a fingerprint protein. Whiskers denote the most extreme observation within 1.5 times of the interquartile range. Data are separated by the populations and are colored based on the population a protein is a predictor for (AK - orange; BH - blue; LB - green; RE - pink).

Two of the four fingerprint proteins identified for the AK population function in osmo- and iono-regulation. G3PMS8 encodes the Na^+^/K^+^-ATPase alpha 1a subunit and G3N4P6 encodes a voltage-gated chloride channel (CLCN2). The higher abundance of these two proteins in the AK population relative to the other populations suggests that active NaCl absorption represents an acute functional need for this population. This conclusion is supported by the location at which these fish were sampled and by their morphotype. The large, fully-plated (trachurus) morphotype of the AK population indicates that they are anadromous and spent most of their life in a marine environment^7^. However, these fish have been sampled in a brackish lagoon at a low salinity that is plasma-hypoosmotic. Therefore, AK fish have an acute need to switch their branchial osmo- and iono-regulation from active NaCl secretion in a marine environment to active NaCl absorption in this plasma-hypoosmotic environment (salinity < 9 g/kg). Such a switch in osmoregulatory gill function is facilitated by up-regulation of the branchial Na^+^/K^+^-ATPase alpha 1a subunit in anadromous teleosts^8^.

The role of CLCN2 for branchial NaCl absorption in euryhaline teleosts is not as well established as that of the Na^+^/K^+^- ATPase alpha 1a subunit. However, there is clearly a need for electrogenic absorption of Cl^−^ along with the absorbed Na^+^ to maintain the electrochemical potential across the gill epithelium. CLCN2 would fulfill this requirement since it carries Cl^−^ in a voltage-dependent manner. However, this function of CLCN2 still has to be validated in gills of teleosts inhabiting FW and low salinity BW. CLCN2 likely functions to close the gap between the significantly greater extent of Cl-that needs to be absorbed across gills of FW fish relative to the amount of 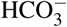 that is excreted by band 3 exchangers.

One of the four proteins has not yet been characterized (G3PH38). It contains weak similarity to the Ig mu heavy chain and may be involved in pattern recognition, but its functional role in teleost gills is unknown. The remaining AK fingerprint protein (G3NUU3) encodes translationally controlled tumor protein 1 (TCTP1). TCTP1 is involved in calcium signaling and the regulation of growth and energy metabolism during stress^9^. Its higher abundance in the AK population likely reflects the osmotic stress encountered by this population when entering low salinity brackish water from a marine environment or being exposed to tidal salinity fluctuations. During such osmotic stress, energy is diverted from growth to osmoregulation, and TCTP1 may be a key protein regulating this process.

A major function that is shared by the two fingerprint proteins for the BH population is their involvement in glycolysis and growth. Fructose-bisphosphate aldolase (FBPA, G3Q1L3) and transgelin (TAGL, G3NRF8) both promote growth processes via stimulation of glycolysis. Transgelin has been investigated as a potential biomarker of accelerated cell proliferation in various cancers^10^. The BH population experiences a very stable environment regarding salinity, temperature, and other key ecological parameters. Thus, energy expenditure for stress responses should be much lower than in the other populations. Moreover, the BH population morphotype (large, fully plated trachurus form) supports the notion that much of the metabolic energy generated is invested in growth processes.

All four fingerprint proteins of the LS population are functionally important during inflammation, suggesting that an inflammatory response is activated in gill tissue of the LS population. Caspase 1 (G3P6H8) is part of the inflammsome complex^11^, beta-2-microglobulin (G3Q6I1) is a common inflammatory protein that has been used as a biomarker of inflammation associated with human diseases^12^, metalloendopeptidases (inlc. choryolysin L-type, G3PN26) are potant targets of anti-inflammatory drugs^13^, and sphingomyelin phosphodiesterases (incl. fatty acid binding proteins, G3PEH9) have been shown to modulate inflammatory responses in many animals^14^. Activation of an inflammatory response in gills of LS fish is supported by their poor growth and significantly smaller size than the AK and BH populations and by their high parasite (*Schistocephalus solidus*) load. Even though fish included in the analysis had been selected for lack of visible parasites, the LS population as a whole was heavily (>50%) visibly parasitized. It is possible that small, non-visible parasites were present or inflammation resulted from fending off parasites in the LS habitat. Alternatively or additionally, inflammation of LS stickleback gills may have resulted from high mercury (Hg) levels in the LS habitat^1^.

The three RE population fingerprint proteins are indicative of preferential protection and/ or repair of basement membranes/ extracellular matrix in gills of RE fish. The RE population represents a margin population existing at their extreme Southern distribution limit. Their southward expansion is limited by the high temperature in brackish environments that minimize energy expenditure for osmoregulation, i.e. environments with a salinity of about 10 g/kg, which is isosmotic to fish plasma. The fingerprint proteins heat shock protein 47 1b (HSP47-1b, G3NT01) and versican B (G3PSB5) indicate significant extracellular matrix damage. HSP47 and versican B counteract damage to the basement membrane and extracellular matrix^15^,^16^. In contrast, DDX3b (G3NZU2) participates in protein quality control, suggesting either a decrease in proteostasis or less stringent protein quality control in the RE population^17^. Stress-induced extracellular matrix damage must be counteracted by a cellular stress response^18^, which is energy-expensive and limited by the amount of energy that can be diverted from growth and other (e.g. osmoregulation) processes. Downregulation of protein turnover (proteostasis) may represent a strategy to minimize energy expenditure in the RE population.

Now that we have identified a fingerprint for the population and explored the biological functions of the selected proteins, we want to investigate the accuracy and robustness of this fingerprint. To get a better understanding of the fingerprint regression model’s performance, we calculate the predicted probability that each sample data point originates from each of the populations. Fig. 3 A) shows this calculation for all four populations separately: The results for predicting AK origin for each of the 96 sample points are in the first column, followed by those testing if the samples originated from BH, LS, and finally RE. The true origin of each sample is encoded in the color (AK, orange; BH, blue; LS, green; RE, pink). For a perfect separation of the four populations, the dots colored orange should have high probabilities in the AK column and low probabilities in the other three columns, and likewise for the other color codes. We note that Fig. 3 A) shows exactly this behavior: a clear separation of the colors between the populations, where the predicted probabilities of the true population is larger than the predicted probabilities of the other populations. Thus, we have confirmed that our LASSO-based 13-fingerprint model is able to correctly assign the origin of the fish with high probabilities.

**Figure 3.**
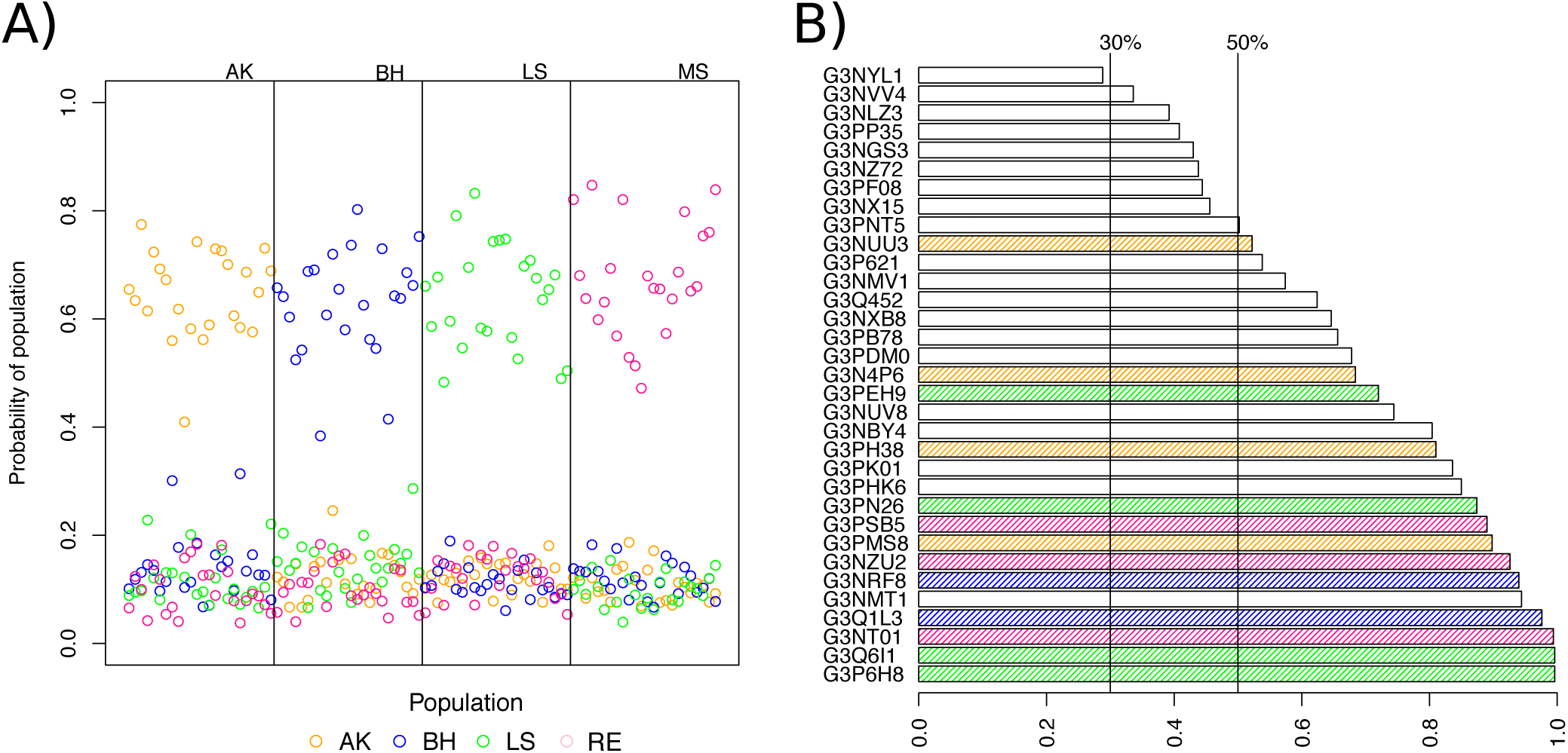
**A)** Probability of originating population for each fish in the data set, assessed for each population. The vertical lines separate which population that is predicted and the vertical placement of the circles shows the probability of this fish (sample) being from a given population. The symbols are colored by the true population membership of the sample. **B)** Most frequently selected proteins in the 500 bootstrap models, where proteins from the 13-protein-model are colored. The vertical lines represents 30% and 50% frequency of inclusion.

As each population only consists of 24 samples, we conduct a robustness analysis of our predictions to ensure that the fingerprint model is not caused by overfitting. Using a bootstrap procedure, we draw 500 bootstrap samples with replacement from the original data (see Methods). For each of the bootstrap samples, we apply the LASSO analysis to fit a new regression model (see Methods section for details). These 500 resulting models are compared in terms of what proteins were selected and their overall prediction accuracy. Ideally we would have the same 13 fingerprint proteins in all bootstrap models, but based on the nature of the LASSO algorithm, some proteins can be replaced by a similarly and highly correlated protein giving an optimal fit to the bootstrapped data, but a different fingerprint. The histogram in Fig. 3 B) shows the most frequently selected proteins in these models. We find that a total of only 25 proteins are selected in more than 50% of the bootstrap models, including all of the proteins from the 13-protein fingerprint. This shows that the fingerprint is quite robust and that there are 12 additional proteins that often appear in the bootstrap fingerprints and could also be interesting proteins to study.

Looking at the prediction accuracy for these bootstrap models, we find that none of the 500 models resulted in any misclassification. When applying the bootstrap models for population prediction of the fish on the original dataset, we find that 96.4% of the bootstrap models resulted in no misclassifications, whereas 3.6% of the models gave rise to one or two misclassifications, and none of the models resulted in more than two misclassifications. Consequently, we have identified a 13-protein fingerprint set that is able to accurately separate and identify the origin of the fish. Our validation of this set shows that the fingerprint proteins are robust, as they are prominent in the 500 similar fingerprints that can be considered accurate fingerprints for the populations.

### Identifying fingerprint proteins for environmental factors

The four stickleback populations have been subject to quite different environmental factors. In addition to identifying proteins specific for the populations, we are also interested in the proteins associated with the fish’s ability to adapt to such different environmental traits; in particular related to the salinity of the water, the temperature of the water and the morphotype of the fish, such as size and lateral plate count. Using the LASSO approach, we generated three models for prediction of these environmental traits, i.e. one model for prediction of habitat salinity, one model for prediction of morphotype, and one model for prediction of habitat temperature. In this way, we identify fingerprint proteins that are specific for the environmental factors and investigate if these fingerprints gives more information than the population fingerprint alone.

#### Salinity fingerprint proteins

We divide the fish into groups based on their habitat’s salinity: Populations AK and RE inhabit brackish water, BH inhabits salt water and LS inhabits FW. Using the LASSO approach to predict these three levels of salinity (see Methods), we find 18 proteins that result in a 98.9% accuracy (meaning one misclassification) in prediction of the habitat salinity each sample inhabit (see Tab. 1 for coefficient estimates). The abundance profiles for these proteins are shown in Fig. 4. Note that the boxes in the figure are separated by population, but marked with the salinity level of the population to highlight the similar pattern for BW fish. Inspecting this figure, we find that high levels of the first three proteins (blue: G3P6H8, G3PN26, G3Q6I1) are specific for FW fish. The next nine proteins (grey) indicate fish from BW where the low levels of seven proteins (G3N8R2, G3NBD5, G3NKU0, G3NUK7, G3PFX9, G3PH93 and Q804E3) and high levels of two proteins (G3NIT4 and G3PYA1) constitute a specific pattern. The proteins in the last row of Fig. 4 indicate the SW fish samples, where the pattern of high levels of these proteins (colored green: G3NMV1, G3NRF8, G3NUV8, G3NX15, G3NXB8 and G3Q1L3) is specific.

**Figure 4.**
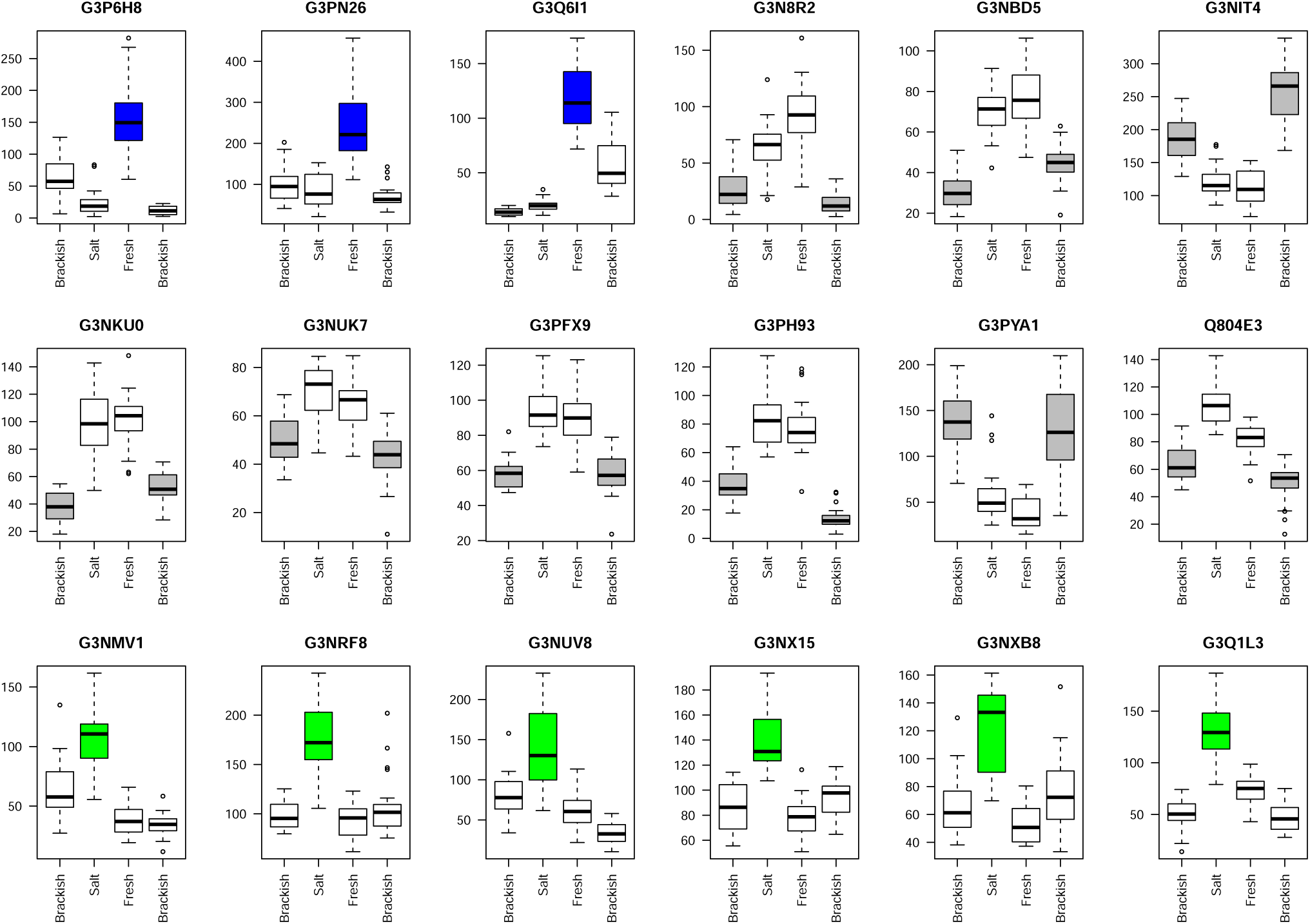
Abundance plot of the salinity-associated fingerprint proteins. The data for each protein are separated into the salinity levels of the four populations and are colored based on the salinity the protein is a predictor for (FW, blue; BW, grey; SW, green). As previously, each box contains the quartiles and the median of the measured abundance levels for a fingerprint protein. Whiskers denote the most extreme observation within 1.5 times the interquartile range.

The three FW specific proteins are identical to fingerprint proteins predicted for the LS population (see above). This is not surprising, as the LS population is the only population representing FW fish in our analysis. Even though there is a strong association of these proteins with FW, it would be wrong to conclude that these proteins are functionally important for osmoregulation. In fact, these proteins are functionally more closely related to other environmental conditions experienced only by the LS population, such as parasitism or Hg contamination. These contexts are specific to LS, but not due to salinity. To truly discern effects of FW on the gill proteome, additional FW populations, ideally from non-contaminated and non-parasitized lakes, would need to be included in the analysis, or laboratory salinity acclimation experiments under controlled (common garden) conditions would have to be conducted to separate such other effects from the salinity effects. These considerations do not lessen the usefulness of the statistical approaches developed here but, instead, they underscore the importance of tailoring the experimental design to the biological question or environmental variable of interest.

The six SW specific proteins include both BH fingerprint proteins (see above). Again, this is not surprising since the BH population was the sole population representing SW fish in our analysis. As for the FW specific proteins, these two SW specific proteins (fructose bisphosphate aldolase and transgelin) do not functionally support adaptation to SW per se, but rather support the associated BH phenotype (large, fast-growing, fully plated). It is possible that transgelin is involved in SW adaptation by promoting gill cell adhesion processes that support active ion secretion under SW conditions. However, such a function of transgelin has not been validated yet.

Three of the other four proteins in this set, succinyl-3-ketoacid coenzyme A transferase (G3NUV8), alpha-enolase (G3NX15), glutathione S-transferase A (GSTA, G3NXB8), are also more related to the high growth morphotype of the BH population than salinity per se. They are metabolic enzymes that promote energy production and growth and (in the case of GSTA) alleviate oxidative stress resulting from high oxidative metabolism. It is possible that the increased abundances of these energy metabolism enzymes also reflect greater needs for osmoregulation in gills, in addition to them supporting the high-growth phenotype. The osmotic gradient between SW and teleost plasma is significantly greater (ca. 25 g/kg) than for FW teleosts (ca. 10 g/kg) or brackish water fish (as low as 0 g/kg)^19^. Thus, a role of these SW specific energy metabolism proteins for energizing the active NaCl secretion processes in gills of SW fish is also plausible. The final SW-associated fingerprint protein, G3NMV1 (carbonic anhydrase), is an enzyme that is central for coordinating two vital physiological functions of fish gills, acid-base regulation and osmoregulation^20^.

In contrast to FW and SW, the effect of brackish water on gill proteomes was interrogated by taking into account two very different populations that have brackish water habitat in common but differ in most other aspects. The two proteins that were elevated under those conditions are structural intermediate filament and cytoskeletal proteins (keratin 18, G3NIT4 and myosin light chain 9, G3PYA1) that stabilize cells. Their high abundance may indicate that gill cells are sturdier in brackish habitats. Most proteins specific to brackish water fish were depleted relative to fish from other salinities. The seven depleted proteins (G3N8R2 Proteasome subunit beta, G3NBD5 Adenylate kinase 2, G3NKU0 Eukaryotic translation initiation factor 4h, G3NUK7 Enoyl CoA hydratase 1, G3PFX9 Electron transfer flavoprotein subunit beta, G3PH93 Peptidylprolyl isomerase, Q804E3 transgelin) are involved in proteostasis (protein turnover) and mitochondrial energy metabolism, which indicates that protein turnover in gill cells of brackish water fish is decreased. A decreased metabolism is consistent with a reduced energetic demand of gill cells for osmoregulation in brackish water.

Looking at the probability plot for habitat salinity shown in Fig. 5 A), we see that the model performs quite well in prediction of habitat salinity. From left, only the true FW fish are assigned a high probability of being a FW fish, and low probabilities for the other salinity levels. Similarly for BW and SW fish (middle and right segment), where the model assigns high probabilities to the fish inhabiting at these salinity levels and low probabilities for the other levels. Thus, we see that the model makes a clear separation between the salinity levels in all except one SW fish, which is in fact the one misclassification of the model. This fish is a true SW fish with a 52% probability of BW and 39% probability of SW.

**Figure 5.**
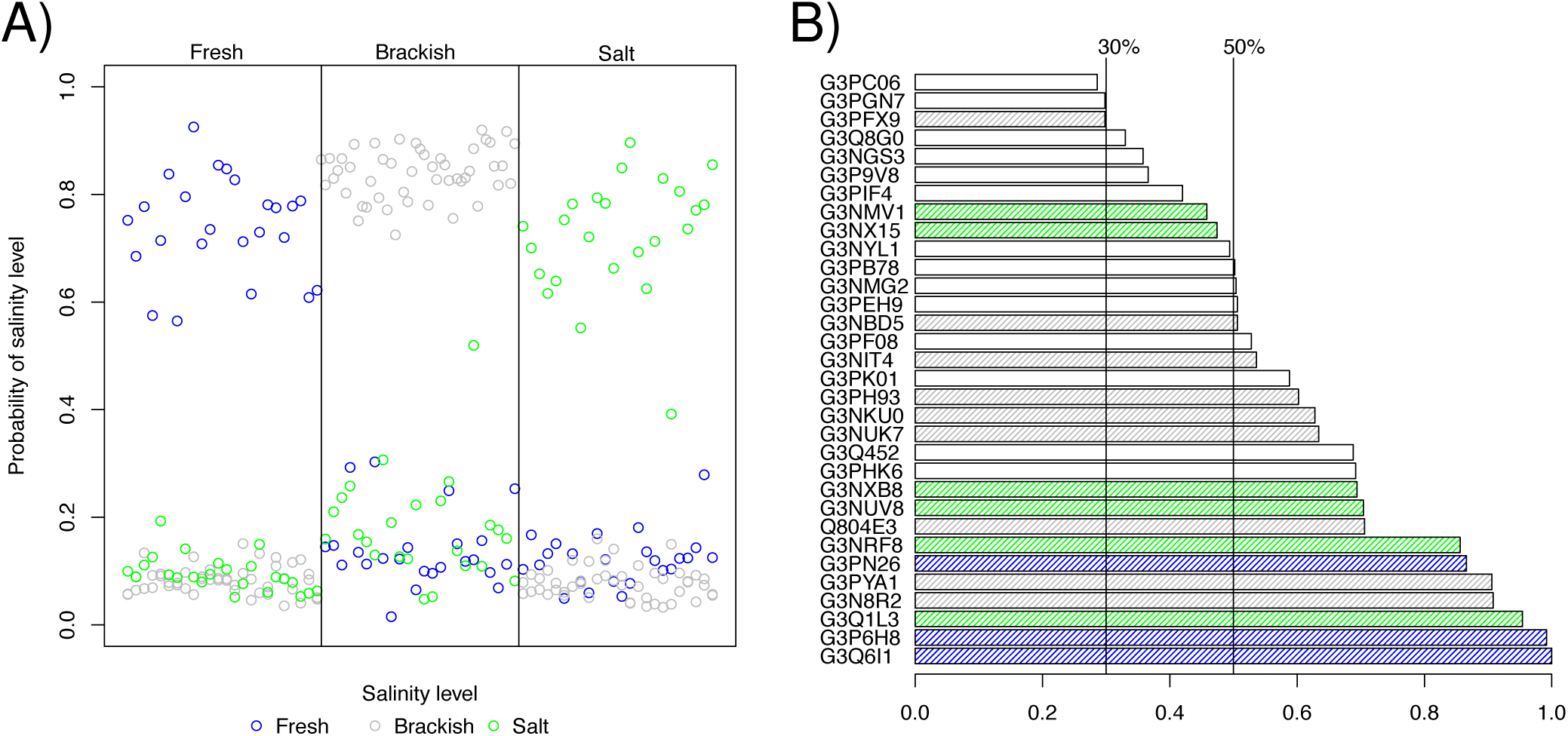
**A)** Probability of originating salinity level for each fish in the data set, assessed for each salinity level. The vertical lines separate which salinity level is predicted and the vertical placement of the circles shows the probability of this fish (sample) being from a population with a given salinity level. The symbols are colored by the true salinity level of the sample point. **B)** Most frequently selected proteins in the 500 bootstrap models, where proteins from the 18-protein-model are colored. The vertical lines represents 30% and 50% frequency of inclusion.

Running a bootstrap procedure as for the population prediction, we find that a total of only 22 proteins are included in more than 50% of the bootstrapped models for prediction of salinity levels. Fig. 5 B) shows that all except three of the salinity fingerprint proteins are among these 22 proteins. It seems that the majority of these fingerprint proteins are very important for prediction, as they appear in most of the bootstrap models, while others can be replaced and the model still performs well in prediction tests. When predicting the habitat salinity of the bootstrapped data, the bootstrap models gave rise to zero misclassifications. On the original data, the bootstrap models did not perform as well for habitat salinity prediction, with only 63.8% of the models resulting in zero misclassifications. However, among the other 36.2% of the models, there are only one or two sample misclassifications for each bootstrap model. Thus, each bootstrap model has an accuracy of 97.9% or higher. Again, this indicates the existence of several accurate “fingerprint” sets of proteins for prediction of habitat salinity, where the inclusion of most of the proteins from our 18-protein fingerprint indicates that our fingerprint is quite robust and accurate.

#### Morphotype fingerprint proteins

To investigate if the data support an identification of protein sets associated with adaptation in morphotype, we apply the LASSO method as follows: The populations AK and BH have the *trachurus* morphotype (large body size, fully plated), while the LS and RE populations have the *leiurus* morphotype (small, low-plated). Using a logistic LASSO model (see Methods), we identified a fingerprint protein set consisting of only four proteins that is reliably capable of separating these morphotypes with zero misclassifications (see Tab. 1 for coefficient estimates). Fig. 6 A) shows that high protein levels of G3NZH1 and G3Q6I1 indicate leiurus and low levels indicate large trachurus morphotypes, while high levels of the proteins G3PLA9 and G3Q6S8 indicate trachurus and low levels indicate leiurus morphotypes. Again, the boxes are separated by population to highlight the similar pattern of the morphotypes.

**Figure 6.**
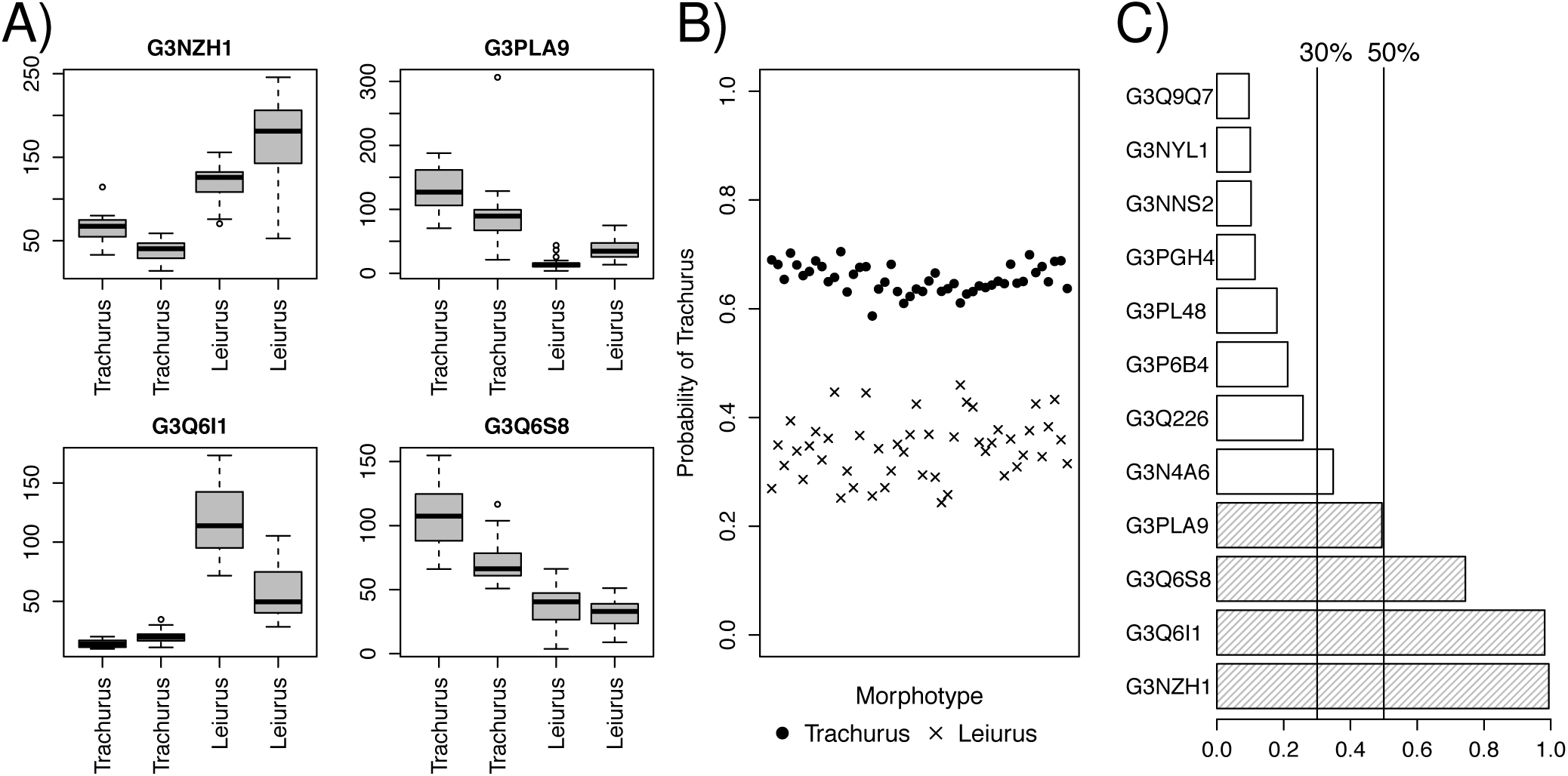
**A)** Abundance plot of the morphotype-associated fingerprint proteins. The data for each protein are separated into the morphotype of the four populations (Trachurus; AK, BH. Leiurus; LS, RE). **B)** Probability of originating the trachurus morphotype for each fish in the data set. The vertical placement of the circles shows the probability of this fish (sample) being from a population with the trachurus morphotype. The symbols mark the true morphotype of the sample point. **C)** Most frequently selected proteins in the 500 bootstrap models, where proteins from the 4-protein-model are striped. The vertical lines represents 30% and 50% frequency of inclusion.

The leiurus-specific proteins (transglutaminase B, G3NZH1, and beta-2-microglobulin, G3Q6I1) are likely indicative of the estuarine and lacustrine environments that represent small bodies of water that both leiurus populations are derived from. In these environments, water quality is poorer than in marine habitats and pollutant concentrations and parasite prevalence are much higher. It is likely that the higher abundances of beta-2-microglobulin and transglutaminase B in gills of leiurus fish functionally reflect these conditions. Both proteins are involved in defenses against pathogen invasion. The role of beta-microglobulin in inflammation and immunity has been mentioned above. In addition, transglutaminase B strengthens the apical side of the gill epithelium by cross-linking cell surface proteins, which increases resilience against pathogen invasion^21^. In addition, both of these proteins could affect bone morphogenesis, which directly relates to morphotype-specific differences. In mammals, beta-2-microglobulin has been shown to form amyloid fibers in bone and tendon under certain conditions^22^. Such fibers may be structurally stabilized by transglutaminase B mediated crosslinking. They may strengthen the epidermis of leiurus fish in the absence of lateral plates. It is possible that these proteins are present in circulating blood and deposited in the epidermis. Since gills are highly vascularized tissues large amounts of blood are included in gill homogenates.

The trachurus-specific proteins (Rab GDP dissociation inhibitor, Rab-GDI, G3Q6S8, and hemoglobin beta, G3PLA9) indicate increased metabolism in gills of AK and BH fish. Hemoglobin beta is present in the blood that perfuses gill tissue. Its increased abundance is associated with a higher oxygen supply to gills of trachurus fish, which supports oxidative metabolism and the larger growth phenotype. Rab-GDI regulates Rab GTPases, which control many cellular signaling processes, including those that are involved in membrane trafficking, cell differentiation and proliferation (growth)^23^.

It is clear from Fig. 6 B) that this set of only four fingerprint proteins is an accurate fingerprint for morphotype, correctly assigning high probabilities of trachurus morphotype for AK and BH fish, and smaller probabilities for LS and RE fish. We assess the robustness of the fingerprint by conducting the bootstrap procedure. Fig. 6 C) shows that the four morphotype fingerprint proteins (striped bars) are included in most of the bootstrap models. G3NZH1 and G3Q6I1 are included in almost 100% of the models, showing that these two are very important for prediction morphotype. G3Q6S8 and G3PLA9 are included in approx. 70% and 50% respectively. This means that these two proteins are sometimes replaced by other proteins in the bootstrap fingerprints (colored white), and still make an accurate prediction of morphotype. Applying these models for prediction of the morphotype, 99% of the models had zero misclassifications and 1% resulted in one misclassification. Predicting the morphotypes of the fish in the original data set, 80.4% of the models had zero misclassifications, while the remaining 19.6% of the models had between one and five misclassifications.

#### Temperature fingerprint proteins

Three of the populations (AK, BH and LS) inhabit cold waters (< 20 °C), while the RE population inhabit warm water (30°C). We use the logistic LASSO model to generate a statistical model for temperature fingerprint proteins. When predicting these two temperature states, we find four proteins that clearly separate the fish samples, as shown in Fig. 7 A) (see Tab. 1 for coefficient estimates). Here, high levels of G3NMT1, G3NT01 and G3PSB5 are specific for fish inhabiting warm water and low levels are specific for cold water, while high levels of G3NZU2 are specific for fish inhabiting cold water and low levels specific for warm water.

**Figure 7.**
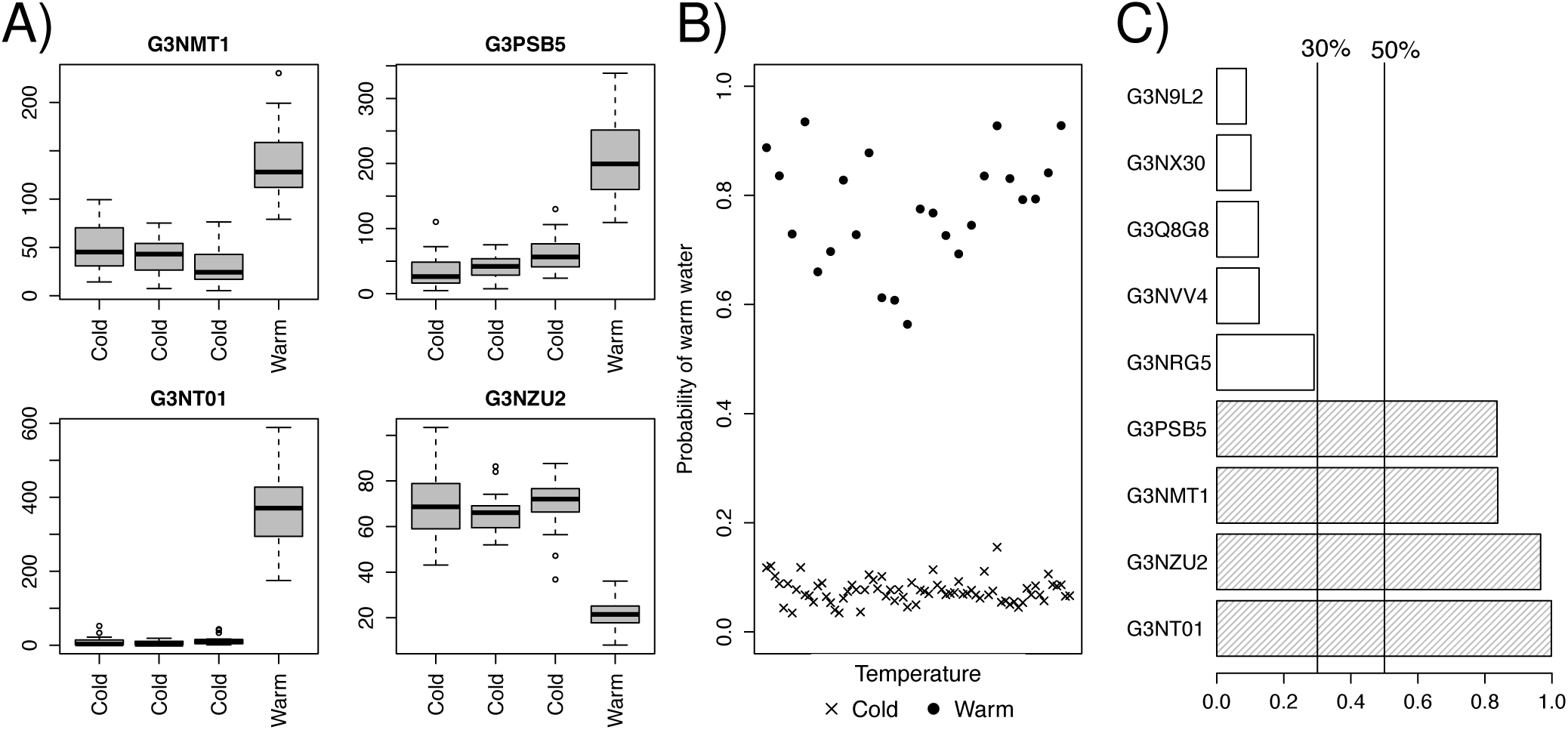
**A)** Abundance plot of the temperature-associated fingerprint proteins. The data for each protein are separated into the temperature of the four populations (Cold; AK, BH, LS. Warm; RE). **B)** Probability of originating warm water for each fish in the data set. The vertical placement of the points shows the probability of this fish (sample) being from a population inhabiting warm water. The symbols mark the true temperature of the sample point. **C)** Most frequently selected proteins in the 500 bootstrap models, where proteins from the 4-protein-model are striped. The vertical lines represents 30% and 50% frequency of inclusion.

The functional significance of three of the four temperature-specific proteins has already been discussed in the context of the RE population-specific proteins (see above). The fourth temperature-specific protein (Septin-type G domain-containing protein, septin 5, G3NMT1) is involved in regulating G protein signaling and functions in the determination of cell polarity and cytoskeleton structure. It is possible that these aspects of gill cell function are affected by temperature effects on basement membrane/ extracellular matrix stability. Thus, higher levels of septin 5 may be required for compensation of such effects.

This model provides a perfect fit to the data with zero misclassifications, and we see from Fig. 7 B) that the model performs very well in separating high-temperature samples from low-temperature ones, assigning very low probabilities for the cold-water fish, while warm-water fish are assigned high probabilities. Using the bootstrap procedure to assess the robustness of this fingerprint, we find that the four temperature fingerprint proteins are included in almost all of the bootstrap models (Fig. 7 C)). This indicates that the abundance pattern of these proteins is highly specific. Looking at the prediction accuracy, we find that 94.4% of the models give a perfect prediction of the bootstrap data, while 5.6% resulted in up to three misclassifications. Predicting the temperature association of the original data, 82.8% of the bootstrap models resulted in a perfect fit, while the remaining 17.2% had up to 7 misclassifications. However, 7 misclassifications for one model gives a prediction accuracy of 92.7%, meaning that all the bootstrap models still perform quite well and that the temperature fingerprint is an accurate and robust fingerprint.

### Identifying proteins with strong differential interactions

Up to now we have identified robust and accurate protein fingerprints for the environments, where the combination of proteins in each fingerprint is specific for the environmental traits. To get a further understanding of how these fish are able to adapt to such different environments, we aim to investigate how the proteins interact and affect each other across the different habitat. A natural limitation with the LASSO is that correlative patterns are not detected, as it aims to find the fewest possible variables to obtain the best fit. Two highly correlated variables are naturally quite similar, and therefore one is sufficient in the LASSO model. To inspect this feature of proteins affecting each other, we take on a system network approach, where we create correlative networks of protein interactions and compare how the interaction differs between the habitats.

Using the Co-expression Differential Network Analysis approach (CoDiNA)^4^, we identify correlative interactions between proteins that differ between the populations. Briefly, the CoDiNA-approach creates a single signed wTO-network for each population, where each protein is represented as a node, and the links represent the wTO-correlative interactions between the proteins (see Methods). Only the strongest wTO-links (adjusted *p*-value < 0.05) are included in each population network and subsequently, the four population networks are merged into one, where a link is present if it is included in one or more of the four networks. The links in the merged CoDiNA-network are classified into three groups based on the type of interactions in the different networks: similar interactions across populations (*α*-links), differential interactions across populations (*β* -links) or specific interactions for one or more population (*γ*-links). Analyzing this network, we are able to identify protein interactions that are similar across the habitats, and more interestingly, protein interactions that have changed between the habitats, which might be evidence for explaining how this fish specie adapt to such different environments.

As we are mostly interested in protein interactions that has changed between the populations, we focus only on the *β* - and *γ*-links representing differential and specific protein interactions across two or three populations. Applying the CoDiNA-approach on our data, we find no statistically significant *β* -links and are therefore left with only *γ*-links to focus on. To highlight the most “extreme” changes between the populations, we consider only the “differential” *γ*-links; protein interactions that are present in two or three populations with opposite sign of the wTO-correlation. Extracting only these links from the CoDiNA-network, we identify a giant component consisting of 117 proteins connected by 139 such links, shown in Fig. 8. The links are colored based on which populations the differential *γ*-link represents. As an example, consider a pink link between two nodes (proteins). These two proteins have either a significantly positive wTO-correlation in the BH population network and a significantly negative link in the RE population network, or vise versa, meaning that these two proteins interact in an opposite way for the fish inhabiting these two populations, while they have no significant interaction for the fish in the AK and LS populations.

**Figure 8.**
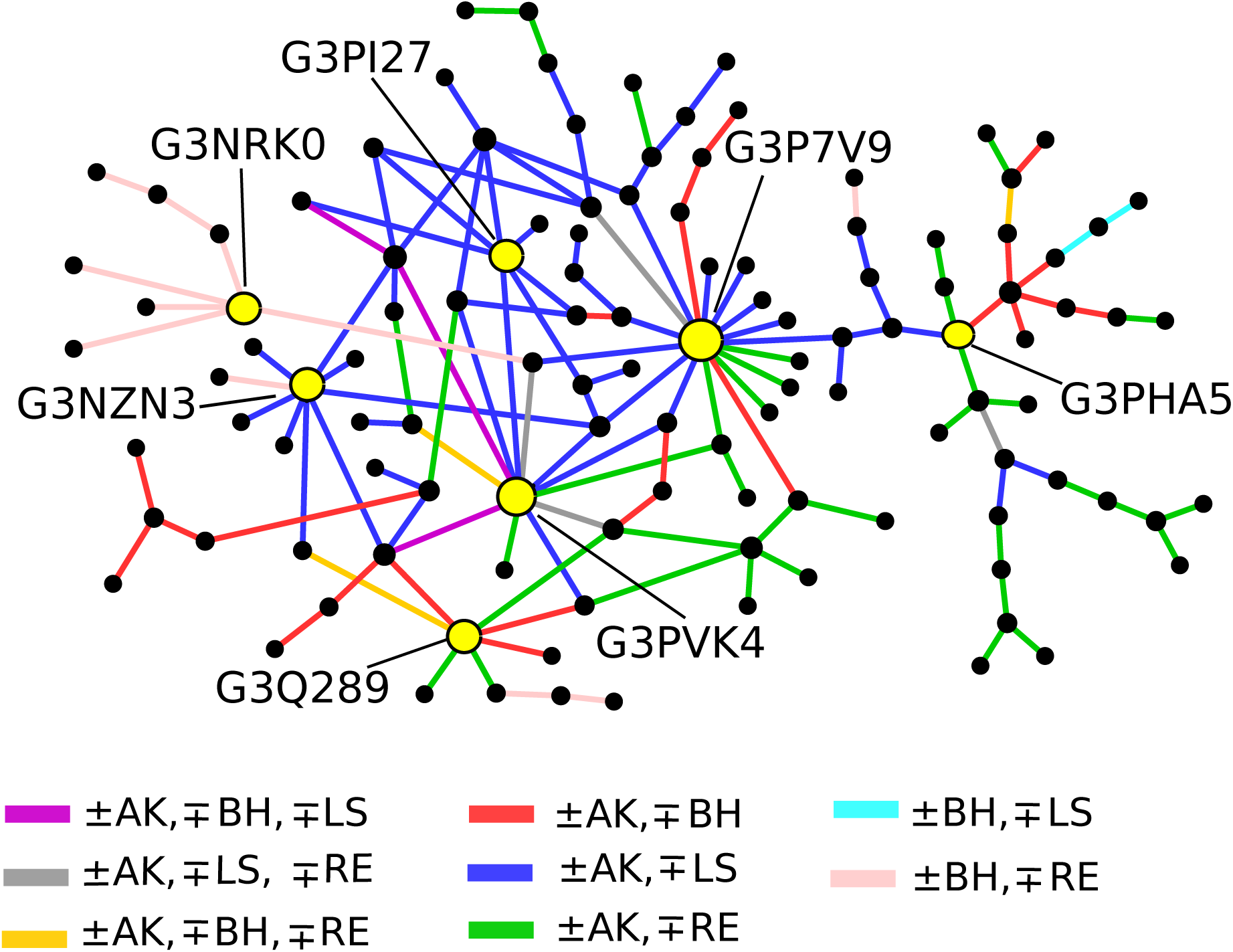
Differential correlation network between four populations: Network of interacting proteins with specific differential interactions in two or three populations. The different link-colors correspond to the patterns of correlation sign-change between the conditions where the link is detected. Seven central nodes are marked in yellow and discussed in more detail in the text.

Inspection of Fig. 8 suggests that links with the same color may cluster in the network, as many of the nodes have links of only one color and branches of nodes are often linked with the same color. We use a permutation test to assess the expected distribution of colors (see Methods) for the nodes as well as the average *H*(*k*)-score (Eq. (3)), where *H*(*k*) represents the average number of different link-colors relative to the node degree *k*.

Fig. 9 shows (A) the color distribution on the nodes and (B) *H*(*k*) for each degree *k* for these simulated networks. The color distribution and *H*(*k*) from our network are marked with blue x-es. It is clear from the figure that our network is dominated by nodes connected to links with a single color, as the number of nodes connected with only one link-color is significantly larger than expected (*p*-value < 10^−4^) and the number of nodes with two or three colors of the links are significantly smaller (*p*-value < 0.05) than expected. With the exception of *k* = 4 (*p* = 0.018), the *H*(*k*) from empirical data is significantly larger than expected (*p*-value < 10^−3^) for all degrees lower than 12.

**Figure 9.**
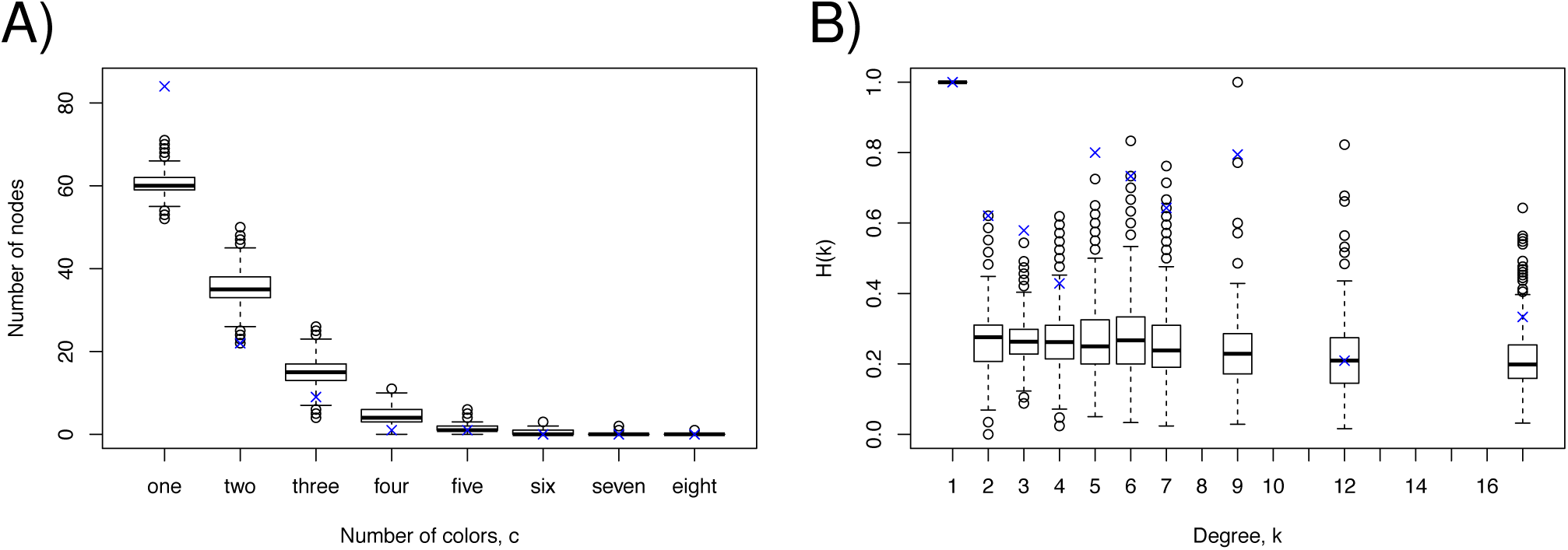
Boxplot of **A)** the simulated color distribution and **B)** *H*(*k*) (see Eq. (3)) in the differential correlation network. Corresponding values from our network is marked by ×.

To develop an understanding of the biological implications of this network, we perform a Fishers Exact over-representation test in PANTHER (protein annotation through evolutionary relationship)^24^ for the proteins in this network. Using the 1490 proteins from our original dataset as reference, the test finds biological functions that are significantly over-represented in the network compared to the reference set. We find that the proteins in our network are significantly (*p*-value < 0.05) enriched for the Hsp90 family charperone protein class (Fold Enrichment 8.51), cell junction (FE 3.44), unfolded protein binding (FE 3.41), actin filament binding (FE 2.74), protein stabilization (FE 12.77), cellular response to heat (FE 6.38), sensory perception of sound (FE 5.32), muscle contraction (FE 3.04), intracellular signal transduction (FE 2.88), cellular component morphogenesis (FE 2.80), protein folding (FE 2.74), actin cytoskeletion organization (FE 2.48), vesicle-mediated transport (FE 2.39) and cellular protein modification (FE 2.09). Several of these functions are related to proteostasis, which is also a key process evident from the fingerprints that predict population, morphotype, salinity and temperature (see above). However, other functions, notably those reflecting the regulation of cytoskeletal organization and morphogenesis are uniquely revealed by the CoDiNA-network. These data suggest that the regulation of proteaostsis, cytoskeletal organization and morphogenesis in stickleback gills is driven by environmental factors that are habitat-specific and/ or by genetic differences between these geographically isolated populations.

We have marked in yellow seven nodes in Fig. 8 that are placed at key branch points, either topological (network hubs) or that connect multiple kinds of differential interactions (connecting links with different colors in Fig. 8). Looking at the biological function of hubs connected to many other proteins with links of one color is interesting, as they have a central role in the network, interacting and affecting many other proteins in a dissimilar matter for a specific population combination. The biological functions of hubs connecting different colors are interesting in the same way, but these proteins have a differential effect on the proteins connected for several population combinations. G3PI27 (heat shock protein 70, HSP70) and G3NZN3 (myosin regulator light chain 9a, MLC9a) are central hubs for the blue links (±AK, ∓LS). These two proteins play major roles in the regulation of proteostasis (HSP70)^25^ and the actin cytoskeleton (MLC9a). The control of actin cytoskeleton by MLC9a is mediated by bone morphogenetic protein (BMP) signaling in other vertebrate cells^26^. Therefore, this pathway may contribute to morphotype differences in stickleback lateral plating and pelvic girdle structure. Moreover, MLC9a also plays a role for regulating tight junctions, which is critical for salinity adaptation^27^. The protein G3NRK0 is a central hub in the pink branch (±BH, ∓RE). This protein encodes ribosomal protein SA, which acts as a laminin receptor suggesting that this node controls gill cell proteostasis and adhesion. The three proteins G3P7V9 (Tumor protein D52), G3Q289 (Gelsolin a) and G3PVK4 (Talin 2a) are hubs with switches, connecting many different colors. These proteins control calcium signaling and cytoskeletal organization. The protein G3PHA5 (Translational (tr)-type guanine nucleotide-binding (G) domain protein) is a switch connecting two branches with different colors to the rest of the network. This protein regulates cell signaling and ribosomal protein synthesis.

A majority of links in the network are colored blue and green, representing different correlation structure of the proteins with these links between population AK and LS (blue) and between AK and RE (green). The environmental factors of population AK and LS differ in salinity (BW vs. FW) and morphotype (trachurus vs. leiurus), while both populations inhabit cold waters. The protein interactions marked with blue could hence evidence specific differential protein interactions between BW and FW and/or trachurus and leiurus morphotype. Similarly, the environmental factors differing for population AK and RE are morphotype (trachurus vs. leiurus) and temperature (cold vs. warm), while both population inhabit BW. The proteins linked by green links could hence be specific differential interaction between trachurus and leiurus morphotype and/or cold and warm temperature.

To understand the biological processes driven by these proteins, we perform a Fishers Exact over-representation test in PANTHER for sets of proteins chained with blue and green links, using the 1490 proteins as a reference set. For the blue set of chained proteins, we find that these proteins are significantly (*p*-value < 0.05) enriched for protein binding (FE 2.16), peptide binding (FE 7.38), heat shock protein binding (FE 11.07), protein binding, bridging (FE 11.07), cytoplasmic vesicle (FE 7.38), chaperonin-containing T-complex (FE 16.6), membrane protein complex (FE 4.15), chaperone protein class (FE 4.33), chaperonin protein class (FE 13.28), peptide metabolic process (FE 16.6), receptor-mediated endocytosis (FE 8.3), membrane invagination (FE 8.3), vesicle budding from membrane (FE 8.3), membrane fusion (FE 6.64) and cellular protein-containing complex assembly (FE 5.86). These enriched functions indicate differences in protein chaperoning by heat shock proteins and alterations in the secretory pathway between the AK and LS populations.

For the green set of chained proteins, we find that these proteins are significantly enriched for the TGF-beta signaling pathway (FE 55.33), Plasminogen activating cascade pathway (FE 41.5), metalloendopeptidase activity (FE 83.0), heme metabolic process (FE > 100), cytoplasmic microtubule organization (FE 83.0), actin filament bundle assembly (FE 27.67), cell junction organization (FE 23.71), protease protein class (FE 5.93), cytoskeletal protein (FE 4.15), actin filament bundle (FE 41.5), focal adhension (FE 33.2) and polymeric cytoskeletal fiber (FE 23.71). This enrichment suggests differential functions of proteolytic systems and different cytoskeletal organization in gills of AK versus RE sticklebacks.

In addition to blue and green links, we see that many links are colored pink and red, representing specific differential protein interactions for population BH and RE and population AK and BH. Population BH and RE inhabit environments differing in all three parameters; salinity (SW vs. BW), temperature (cold vs. warm) and morphotypes (trachurus vs. leiurus), while population AK and BH only differs in the salinity habitat (BW vs. SW). Pink links could hence be evidence for specific differential protein interactions for all three parameters, while red links could be evidence for specific differential protein interactions for BW and SW. The Fishers Exact over-representation test for chained proteins linked with these colors shows that the proteins chained with red colors are enriched for hydrolase activity (FE 4.16), inositol phosphate phosphatase activity (FE > 100), proteasome core complex, alpha-subunit complex (FE 30.49), cell surface (FE 23.71), perinuclear region of cytoplasm (FE 53.36), actin and actin related protein class (FE 26.68), translation elongation factor protein class (FE 23.71), Hsp90 family chaperone protein class (FE 71.14), organophosphate catabolic process (FE > 100), protein stabilization (FE > 100), cellular carbohydrate metabolic process (FE 71.14), alcohol metabolic process (FE 42.69), small molecule catabolic process (FE 35.57), cellular response to heat (FE 35.57), intracellular protein transport (FE 10.41) and vesicle-mediated transport (FE 8.89). These data indicate that proteostasis, the secretory pathway and inositol metabolism represent the major functional differences between BH and AK populations.

For the pink set of chained proteins, we find that these proteins are significantly enriched for the p53 pathway (FE 31.13), the FGF signaling pathway (FE 23.34), protein phosphatase regulator activity (FE > 100), protein phosphatase binding (FE 62.25), protein serine/threonine phosphatase activity (FE 46.69), signaling receptor activity (FE 31.13), heme binding (FE 23.34), 90S preribosome (FE 93.38), t-UTP complex (FE 46.69), cell junction (FE 21.55), HMG box transcription factor (FE 46.69), protein phosphatase (FE 37.35), chromatin/chromatin-binding protein (FE 31.13), cleavage involved in rRNA processing (FE > 100), maturation of 5.8S rRNA from trisistronic rRNA transcprits (FE < 100), rRNA-containing ribonucleoprotein complex export from nucleus (FE > 100), RNA 3’-end processing (FE 93.38), positive regulation of autophagy (FE 62.25), cell cemotaxis (FE 62.25), positive regulation of cytokine secration (FE 62.25), interleukine-6 production (FE 62.25), protein secration (FE 62.25), positive regulation of ERK1 and ERK2 cascade (FE 62.25), NIK/NF-kappaB signaling (FE 62.25), regulation of signaling receptor activity (FE 62.25), autophagy (FE 62.25), activation of innate immune response (FE 62.25), sensory perception of sound (FE 46.69), maturation of SSU-rRNA from tricistronic rRNA transcript (FE 46.69), positive regulation of NF-kappaB transcription factor activity (FE 37.35), MAPK cascade (FE 37.35), chromatin remodeling (FE 37.35), innate immune response (FE 31.13), drug transport (FE 26.68), DNA recombination (FE 26.68), positive regulation of transcription by RNA polymerase II (FE 23.34), cellular component morphogenesis (FE 13.66) and muscle contraction (FE 13.34). Therefore, the main functional differences captured in the gill proteome of BH versus RE sticklebacks relate to intracellular signaling pathways that control many essential cell functions, including the cellular stress response (CSR), cell and epithelial morphogenesis, and immunity. These far-reaching functional differences are consistent with differences between BH and RE sticklebacks in salinity, temperature, and morphotype (in addition to population origin). All other population comparisons differ in only one or two (but not all three) of these parameters.

## Discussion

In this paper, we have used data-driven approaches to derive proteomic fingerprints and proteomic network structures for environmental traits. In contrast to genomic fingerprinting, proteomic fingerprinting captures the dynamic nature of population responses to specific environmental contexts. Because these contexts change dynamically, proteomic fingerprinting is not as suitable for absolute determination of population association of a particular fish as is genomic fingerprinting. Such association will only be appropriate if fish that are at the same life history stage and have been exposed to similar environmental contexts. However, a fundamental advantage of proteomic fingerprinting over genomic fingerprinting is its potential for bioindication.

Because of the context-specific nature of proteomic fingerprints, they reflect prior exposures to specific environments and conditions. Bioindication based on a single or few arbitrarily chosen indicator proteins (biomarkers) has proven problematic and was not very successful by and large in the past. The reasons for these difficulties were often due to choosing common stress-proteins as biomarkers that respond strongly but non-specifically to many different types of stress (e.g. HSP70, HSP90, antioxidant enzymes, etc.). By experimentally deriving fingerprint sets of proteins that specifically distinguish contexts of interest it is possible to develop more discriminatory and complex biomarkers.

It will be interesting to investigate how the set of proteins for discrimination will grow or shrink and whether some of these 13 proteins from the population fingerprint are retained or lost as additional populations are added to the comparison. Moreover, it will be of interest to correlate ecotypes and morphotypes with specific sets of fingerprint proteins and statistically assess whether certain fingerprint sets correspond to specific ecotype/ morphotype features independent of population and/or whether there are some true population-discriminatory proteins left if many populations are analyzed in this way. The latter scenario would indicate genetic differences that affect protein expression levels in different populations. Furthermore, using our approach specific fingerprint protein sets could be associated with particular environmental contexts (e.g. salinity, temperature, parasitism and combinations thereof) by isolating the parameters (or combinations of parameters) of interest in laboratory acclimation experiments under controlled (common garden) conditions. Such studies are now possible by applying the system-level tools developed in the current study.

One way to increase discriminatory power of our approach is to include proteomes of additional tissues in the analysis. This will increase the complexity of data roughly by multiplying the number of proteins per tissue (ca. 1,500) by the number of tissues included in the analysis. The cost of data generation would be significantly greater by including multiple tissues and from a statistical perspective the number of replicates should also be greater. In the current analysis, only 24 samples are used to identify fingerprint sets from ca. 1,500 proteins. Even though our results seems to be quite robust with such few samples, one would prefer more samples to get more certain results. Adding multiple tissues to the analysis, we would require even more samples to obtain trustful results. It might be prudent to investigate to what extent discriminatory power gains from including additional tissues and to test which tissues are most informative for discerning genetic differences between populations and predicting prior environmental exposures within and between populations.

Based on the results produced in this paper, we can not conclude that the findings are purely based on environmental exposures. There could be genetic differences causing the differences we observe, as the genetic and exposure differences are both bunched together, which makes it more difficult to interpret the causes for the differences. An alternative and interesting aim in a future experimental design could be to disentangle genetic from exposure differences, to asses and isolate both the genetic effect and the environmental effect.

As briefly discussed in the result section, a drawback with the LASSO method is that it does not include highly correlated variables in the set of selected variables. For our analysis, this means that the fingerprints could lack several important proteins that distinguish the environmental traits, as they could be highly correlated with proteins already included in the fingerprint. For prediction quality, adding the highly correlated proteins to the model will not contribute to increasing the model accuracy, but for model-inference it could be interesting to see how these correlated proteins could explain the environmental differences. As for now, to the best of the authors knowledge, there exists no solid frameworks for handling correlative patterns with LASSO. If such a framework would be implemented in the future, it could be interesting to see how the fingerprints change and how other proteins could contribute to the models. In this paper, we have worked our way around this drawback by running a bootstrap procedure to see if the fingerprints are missing some informative proteins, and by performing a network analysis where we explicitly consider the correlation structure of the proteins. In this way we study both the most accurate fingerprints in terms of predicting the environmental traits and the protein interactions that differs the most between the environments.

## Methods

### Stickleback gill proteomics data

In this analysis, we used data previously published^1^. Briefly described, this dataset consists of measurements on three-spine sticklebacks (*Gasterosteus aculeatus*) that were collected from populations that inhabit environments of different temperatures (15°C and 30°C) and salinities (fresh water, brackish water, sea water). The sampled fish also represent different morphotypes (trachurus = large size, fully plated; leiurus = small size, low plated). The preparation of gill samples for proteomics and the online LC-MS analysis by data-independent acquisition (DIA) quantitative proteomics have also been previously described^1^. The following proteomics data sets were used for the analyses developed in this study: 1506 protein abundance measurements from 24 fishes sampled from Westchester Lagoon, Alaska (AK; cold brackish water, trachurus morphotype), Bodega Harbor, California (BH; cold marine water, trachurus morphotype), Lake Solano, California (LS; cold fresh water, leiurus morphotype) and Laguna de la Bocana del Rosario, Mexico (RE; warm brackish water, leiurus morphotype), in total 96 samples for each protein. All DIA data and metadata including the *G. aculeatus* gill-specific assay library were derived from the openly accessible Panorama Public proteomics repository (https://panoramaweb.org/labkey/JoLi-01.url) as previously described^1^.

### Imputation

Among the 1,506 protein abundance measurements, 552 of the proteins have one or more measurements of zeros. A zero measurement can either represent that there is no abundance of this proteins for the specific fish or a missing value. Inspecting the data for patterns of true missing values, we removed 16 proteins that had either zero measurements for many fish in the same population or too many (> 30) zero measurements across the populations. For the remaining proteins with more than 16 zero measurements, the abundance distributions for each protein are quite dense without large tails, and imputation can be applied.

Since there is only 24 samples per population, an imputation method based on drawing randomly from these would be vulnerable to bias. Instead, we use the statistically more robust approach of simulating a distribution based on quantiles as follows for the imputation of all the remaining proteins with zero measurements: for each protein with zero measurements: 1) consider all non-zero measurements for this protein in one population at a time and simulate 100 values from a uniform distribution between each 5% quantile interval of the distribution for these measurements. 2) Replace zero measurement with randomly drawn values from the simulated distribution. 3) Repeat the two steps for each of the remaining populations containing zero-measurements for this protein. This procedure leaves us with 96 samples (fish) all with abundance measurements of 1490 proteins.

### Variable selection using LASSO

To identify sets of “fingerprint” proteins that are specific for the environment of the fish we apply the LASSO^2^, a well-known statistical method for subset selection and prediction in a high dimensional regression setting^28^. Treating the protein abundance as covariates and the environmental traits (population, salinity, morphotype or temperature) as the outcome in a regression model, we are in a high dimensional setting where the number of samples, *n*, are less than the number of covariates, *p*, (*p* ≫ *n*), and a simple regression model can not be used. The LASSO method, however, is able to use all the proteins as covariates in the model, and by adding a penalty on the number of coefficients, it shrinks the coefficients that are less important for the prediction of the trait to zero. In this way, LASSO identifies only a small set of variables that are most specific for the trait.

Treating the outcomes *Y*_1_, …,*Y*_*n*_ as independent random variables from a probability density function *f*(*y*_*i*_, *η*(*β*_0_, *β*)), we assume that the outcomes depend on some function of the linear predictor, *η* = *β*_0_ + *x*_1_*β*_1_ + … + *x*_*p*_*β*_*p*_, for the *p* coefficients (proteins). Adding a penalty term the the negative log likelihood function, −*l*(*y*_*i*_, *η*(*β*_0_, *β*)), the LASSO puts a constraint to the number of coefficients allowed in the model. Minimizing this objective function with respect to the *p* coefficients,

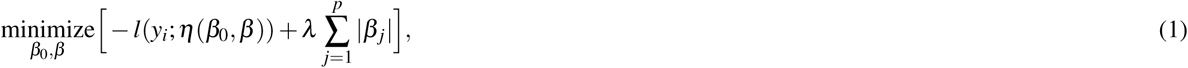

the coefficients are shrunken towards zero, and only a smaller set of non-zero coefficients are left in the model, giving the optimal fit for the given tuning parameter *λ*. The tuning parameter is determined using k-fold cross validation, where the data is randomly divided into 10 folds. The model is trained (minimizing the objective function in Eq. (1)) with subsequently 9 folds and tested on the remaining fold, for a range of *λ* values. We select the value of *λ* with the smallest mean misclassification error, and the final model is given by the estimated coefficients giving the optimal fit with this *λ* value.

#### Multinomial LASSO

When modelling the population or the salinity of the fish, the outcome is a factor variable with more than two levels (population; four levels, salinity; three levels). For these models, a multinomial LASSO is applied^6^, where the outcome depends on *K* linear predictors, *η*_*k*_ = *β*_0_ + *x*_1_*β*_1,*k*_ + … + *x*_*p*_*β*_*p,k*_, one for each level. In this case, each fish is assigned a probability for each level *k* ∈ {1,.., *K*},

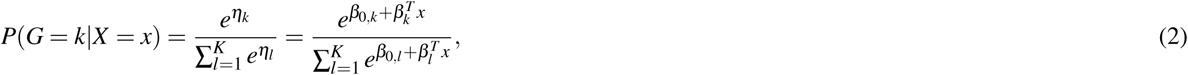

where *x* is the matrix of observed protein abundance and *β*_*k*_ is the coefficient vector for level *k*. Minimizing the multinomial negative log likelihood subject to the penalty factor,

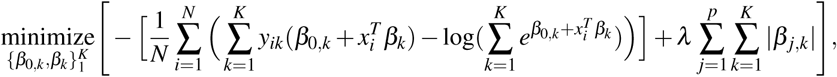

and performing a 10-fold cross validation, each level of the outcome will have a specific linear predictor, where most of the estimated coefficients are zero. Note that positive values of the estimated coefficients contribute to higher probability (evidence for) this level, while a negative estimated value contributes to lowering the probability (evidence agains) this level. Using the multinomial family with the ungrouped version of the penalty factor in the R-package “glmnet”^29^, we ensure that the non-zero coefficients do not need to be non-zero for all levels. When testing the model, each fish is assigned a probability for each level according to Eq.(2) with the estimated coefficient values. The fish is classified to the level with the highest probability.

#### Logistic LASSO

For modelling the size and temperature of the water, the outcome is a factor variable with only two levels and a logistic LASSO is applied^6^. The procedure is very similar as for a multinomial model, but now the outcomes only depend on one linear predictor, *η* = *β*_0_ + *x*_1_*β*_1_ + … + *x*_*p*_*β*_*p*_, and each fish will only be assigned a probability for success (trachurus morphotype or warm water),

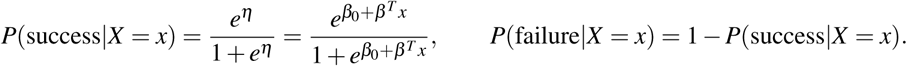

Hence, a probability of success close to 1 indicate high probability for trachurus morphotype/ warm water, while a probability close to 0 indicate leiurus morphotype/ cold water. Minimizing the logistic negative log likelihood subject to the penalty factor,

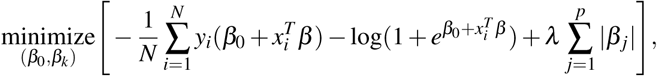

and performing a 10-fold cross validation, the LASSO identifies a set of non-zero coefficients that important for determining the morphotype/temperature. Here, positive values of the estimated coefficients contribute to the evidence for trachurus morphotype/ warm water, while negative values contribute to the evidence for leiurus morphotype/ cold water. This model is fitted using the binomial family in “glmnet”^29^.

#### LASSO application

We estimate the models using the above described statistical framework. For the population model as an example, we estimate the coefficients *β* that give the optimal fit using

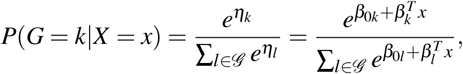

where *k* ∈ *ℊ* = {*AK, BH, LS, RE*} are the four different population origins (levels), and *η*_*k*_ is the linear predictor for each level. After execution of the LASSO algorithm, we identify the following linear predictors as optimal:

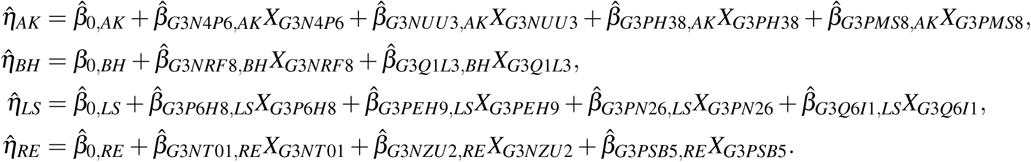

The estimated coefficients (fingerprints) for all of the statistical models are shown in table 1.

**Table 1.**
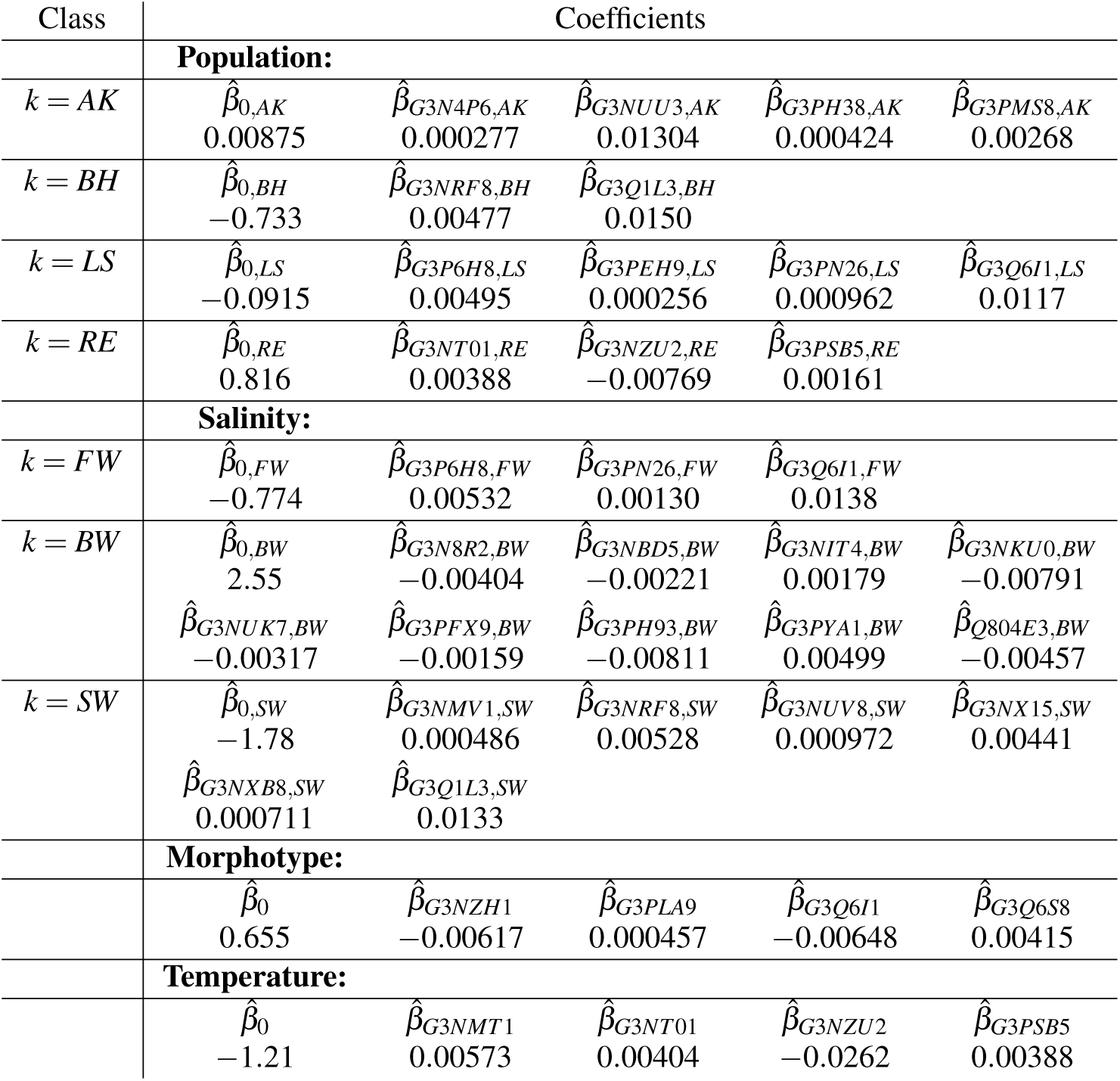
The fingerprint statistical models using LASSO for the four tests: Population, Salinity, Morphotype and Temperature. For population model, AK, BH, LS, RE correspond to the four separate population sample origins. For the salinity model, FW, BW and SW correspond to the three salinity levels found in the population environments. The morphotype and temperature models are logistic models with only one set of coefficients.

| Class | Coefficients |  |  |  |  |
| --- | --- | --- | --- | --- | --- |
|  | **Population:** |  |  |  |  |
| $k = AK$ | $\hat{\beta}_{0,AK}$<br>0.00875 | $\hat{\beta}_{G3N4P6,AK}$<br>0.000277 | $\hat{\beta}_{G3NUU3,AK}$<br>0.01304 | $\hat{\beta}_{G3PH38,AK}$<br>0.000424 | $\hat{\beta}_{G3PMS8,AK}$<br>0.00268 |
| $k = BH$ | $\hat{\beta}_{0,BH}$<br>-0.733 | $\hat{\beta}_{G3NRF8,BH}$<br>0.00477 | $\hat{\beta}_{G3Q1L3,BH}$<br>0.0150 | | |
| $k = LS$ | $\hat{\beta}_{0,LS}$<br>-0.0915 | $\hat{\beta}_{G3P6H8,LS}$<br>0.00495 | $\hat{\beta}_{G3PEH9,LS}$<br>0.000256 | $\hat{\beta}_{G3PN26,LS}$<br>0.000962 | $\hat{\beta}_{G3Q6I1,LS}$<br>0.0117 |
| $k = RE$ | $\hat{\beta}_{0,RE}$<br>0.816 | $\hat{\beta}_{G3NT01,RE}$<br>0.00388 | $\hat{\beta}_{G3NZU2,RE}$<br>-0.00769 | $\hat{\beta}_{G3PSB5,RE}$<br>0.00161 | |
|  | **Salinity:** |  |  |  |  |
| $k = FW$ | $\hat{\beta}_{0,FW}$<br>-0.774 | $\hat{\beta}_{G3P6H8,FW}$<br>0.00532 | $\hat{\beta}_{G3PN26,FW}$<br>0.00130 | $\hat{\beta}_{G3Q6I1,FW}$<br>0.0138 | |
| $k = BW$ | $\hat{\beta}_{0,BW}$<br>2.55 | $\hat{\beta}_{G3N8R2,BW}$<br>-0.00404 | $\hat{\beta}_{G3NBD5,BW}$<br>-0.00221 | $\hat{\beta}_{G3NIT4,BW}$<br>0.00179 | $\hat{\beta}_{G3NKU0,BW}$<br>-0.00791 |
| | $\hat{\beta}_{G3NUK7,BW}$<br>-0.00317 | $\hat{\beta}_{G3PFX9,BW}$<br>-0.00159 | $\hat{\beta}_{G3PH93,BW}$<br>-0.00811 | $\hat{\beta}_{G3PYA1,BW}$<br>0.00499 | $\hat{\beta}_{Q804E3,BW}$<br>-0.00457 |
| $k = SW$ | $\hat{\beta}_{0,SW}$<br>-1.78 | $\hat{\beta}_{G3NMV1,SW}$<br>0.000486 | $\hat{\beta}_{G3NRF8,SW}$<br>0.00528 | $\hat{\beta}_{G3NUV8,SW}$<br>0.000972 | $\hat{\beta}_{G3NX15,SW}$<br>0.00441 |
| | $\hat{\beta}_{G3NXB8,SW}$<br>0.000711 | $\hat{\beta}_{G3Q1L3,SW}$<br>0.0133 | | | |
|  | **Morphotype:** |  |  |  |  |
| | $\hat{\beta}_0$<br>0.655 | $\hat{\beta}_{G3NZH1}$<br>-0.00617 | $\hat{\beta}_{G3PLA9}$<br>0.000457 | $\hat{\beta}_{G3Q6I1}$<br>-0.00648 | $\hat{\beta}_{G3Q6S8}$<br>0.00415 |
|  | **Temperature:** |  |  |  |  |
| | $\hat{\beta}_0$<br>-1.21 | $\hat{\beta}_{G3NMT1}$<br>0.00573 | $\hat{\beta}_{G3NT01}$<br>0.00404 | $\hat{\beta}_{G3NZU2}$<br>-0.0262 | $\hat{\beta}_{G3PSB5}$<br>0.00388 |

#### Fingerprint data validation

A well-known method for validation of statistical models, such as those arising from LASSO analyses, is bootstrapping^6^, 30. Here, a bootstrap sample consisting of 96 data points is drawn with replacement from the original sample, and we repeat the process *B* = 500 times. For each of the *B* bootstrap samples, a LASSO model is fitted, resulting in *B* regression models with the best fit for each of the *B* samples. In order to determine the most important proteins, we assess the frequency by which each protein is selected in the *B* bootstrap models. We subsequently validate the bootstrap models by identifying the number of misclassifications each bootstrap model produces when applying the *B* models on both their respective bootstrap data samples and on the original data.

In this way, we validate the robustness of the assessed fingerprint; if the fingerprint proteins appear in most of the bootstrap models, and if these bootstrap models, consisting of many of the fingerprint proteins, are accurate in prediction we have a robust and accurate fingerprint.

### Network approach to identify proteins with opposite functions

Considering each protein as a node, we apply the differential network approach CoDiNA^4^ to detect specific, common or differential links of interacting proteins among the four populations. The CoDiNA approach takes as input a single network for each condition to be compared. For each of the populations, we generate a separate network using the “wTO”^31^ R-package in the following way: First, a pairwise correlation adjacency matrix is calculated. Second, the correlation values are used to calculate the weighted Topological Overlap (wTO)^28^, 32 between each protein pair (*i, j*),:

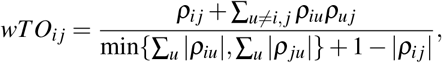

where *ρ*_*i j*_ = corr(*i, j*) is the pairwise Pearson correlation between proteins *i* and *j*. Using the wTO-scores as link measures in networks has been shown to be a biological meaningful measure of interactions^32^–34, where all nodes are connected to every other node, but the strengths of the links differ.

Third, to ensure that only the strongest links are included, a filtering step based on the significance of the link scores are applied^31^: A null hypothesis of the link score being equal to zero is tested against the alternative hypothesis that the link scores are different than zero. Using a bootstrap procedure, the individual data are resampled to approximate the weights’ empirical distribution and probabilities of the observed weights being sufficiently different from zero are calculated. The *p*-values are adjusted for multiple testing with the Bejamini-Hochberg procedure^35^, controlling the false discovery rate. Only links with adjusted *p*-value less than 0.05 are included in the population networks that are input for CoDiNA.

Finally, using the CoDiNA R-package, the four population networks are merged into one differential network where links are classified into three categories based on the type of interaction in each network: An *α*-link represents similar interactions, i.e. a correlative interaction that is present in all four population networks with the same sign of the wTO-score, a *β* -link represents differential interactions, i.e. an interaction that is present in all four networks with opposite signs of the wTO-scores (opposite for at least one population), and a *γ*-link represents specific interactions, i.e, interactions that are present in one, two or three of the networks (possibly with opposite sign of wTO-score), but not being present in the remaining population(s). Considering only nodes connected with *β* - and/or *γ*-links, we identify sets of nodes that interact differently for the specific environments. We note that the largest hub of the network has *k* = 17 nearest neighbors.

#### Assessment of link-color distribution

We use a random permutation test to assess the empirical distribution of link-colors in the differential network by simulating 10^4^ networks with the same structure, while randomly assigning the colors of the links and following the link color distribution. In this way, we keep the network properties unchanged, such as node degree, number of links and distribution of colors among the links, but we randomize the placement of the link colors.

For each network, we calculate two properties: (1) the color histogram, which is the number of nodes that have links of *n* different colors out of the maximum *C* = 8 colors, and (2) the mean *H*(*k*)-index for nodes of degree *k*, where *H*(*k*) is defined as

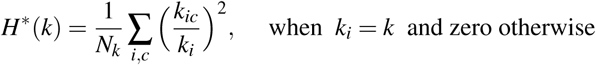

with *k*_*ic*_ is the number of links with the specific color *c* connected to node *i, k*_*i*_ is the total number of nearest neighbors (degree) of node *i*, and *N*_*k*_ is the total number of nodes in the network with degree *k* ^36^. To get a scoring function that takes values between zero and unity, we use

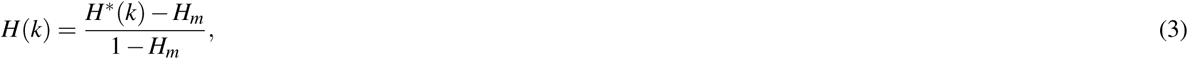

where *H*_*m*_ is the minimum value of *H*^*^(*k*) with the following values

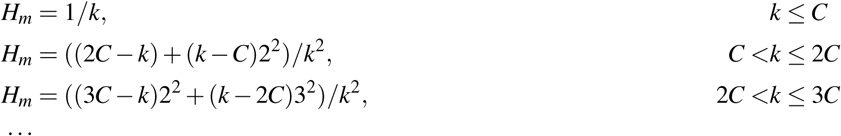

A value of *H*(*k*) = 1 means that all nodes with degree *k* only have links with a single color (not necessarily the same color), whereas a value of *H*(*k*) = 0 means that the link colors for all nodes with degree *k* are maximally different.

## Acknowledgements

M.H. and E.A. would like to thank K. G. Jebsen Foundation grant SKGJ-MED-015. DK was supported by NSF grant IOS-1656371.

## Author contributions statement

D.K. and E.A. conceived the project. M.H. and E.A. developed the project. M.H. conducted the computational analysis. M.H. and E.A. analyzed the results. M.H., D.K., and E.A. wrote the manuscript. All authors approve of the submission, have read and approved the final manuscript.

## References

1. Li, J., Levitan, B., Gomez-Jimenez, S. & Kültz, D. Development of a gill assay library for ecological proteomics of threespine sticklebacks (gasterosteus aculeatus). Mol. Cell Proteomics 17, 2146–2163 (2018).

2. Tibshirani, R. Regression shrinkage and selection via the lasso. J. Royal Stat. Soc. Ser. B (Methodological) 28 (1), 267–288 (1996).

3. Kültz, D. et al. Population-specific renal proteomes of marine and freshwater three-spined sticklebacks. J. Proteomics 135, 112–131 (2016).

4. Gysi, D. M., Fragoso, T. M., Buskamp, V., Almaas, E. & Nowick, K. Comparing multiple networks using the co-expression differential network analysis (codina). ArXiv: Stat.CO 1802.00828 (2018).

5. Pearson, K. LIII. On lines and planes of closest fit to systems of points in space. The London, Edinburgh, Dublin Philos. Mag. J. Sci. 2, 559–572 (1901).

6. Hastie, T., Tibshirani, R. & Wainwright, M. Statistical Learning with Sparsity: The Lasso and Generalizations (Chapman & Hall, 2015).

7. Paepke, H. Die Stichlinge (Neue Brehm-Bücherei, Lutherstadt Wittenberg, 1996).

8. McCormick, S. D., Regish, A. M. & Christensen, A. K. Distinct freshwater and seawater isoforms of na+/k+-atpase in gill chloride cells of atlantic salmon. Exp. Biol 212, 3994–4001 (2009).

9. Kobayashi, D. et al. Identification of a specific translational machinery via tctp-ef1a2 interaction regulating nf1-associated tumor growth by affinity purification and data-independent mass spectrometry acquisition (ap-dia). Mol Cell Proteomics 18, 245–262 (2019).

10. Dvorakova, M., Nenutil, R. & Bouchal, P. Transgelins, cytoskeletal proteins implicated in different aspects of cancer development. Expert. Rev. Proteomics (2014).

11. Angosto, D. et al. Evolution of inflammasome functions in vertebrates: Inflammasome and caspase-1 trigger fish macrophage cell death but are dispensable for the processing of il-1beta. Innate Immun 18, 815–824 (2012).

12. Argyropoulos, C. P. et al. Rediscovering beta-2 microglobulin as a biomarker across the spectrum of kidney diseases. Front Med (Lausanne) 4, 73 (2017).

13. Gearing, A. J. H., Adams, S. E., Clements, J. C. & Miller, K. M. Matrix metalloproteinases in neuro-inflammatory disease, 85–98 (Birkhäuser Basel, Basel, 1999).

14. Hui, X. et al. Adipocyte fatty acid-binding protein modulates inflammatory responses in macrophages through a positive feedback loop involving c-jun nh2-terminal kinases and activator protein-1. J Biol Chem 285, 10273–10280 (2010).

15. Wight, T. N. Provisional matrix: A role for versican and hyaluronan. Matrix Biol 60–61, 38–56 (2017).

16. Verrico, A. K. & Moore, J. V. Expression of the collagen-related heat shock protein hsp47 in fibroblasts treated with hyperthermia or photodynamic therapy. Br. J. Cancer 76, 719–724 (1997).

17. Padmanabhan, P. K. et al. Ddx3 dead-box rna helicase plays a central role in mitochondrial protein quality control in leishmania. Cell Death Dis 7, e2406 (2016).

18. Kültz, D. Molecular and evolutionary basis of the cellular stress response. Annu. Rev Physiol 67, 225–57 (2005).

19. Kültz, D. Physiological mechanisms used by fish to cope with salinity stress. J. Exp. Biol 218, 1907–1914 (2015).

20. Gillmour, K. M. New insights into the many functions of carbonic anhydrase in fish gills. Respir Physiol Neurobiol 184, 223–230 (2012).

21. Cruz, S. I. R., Phillips, M. A., Kültz, D. & Rice, R. H. Tgm1-like transglutaminases in tilapia (oreochromis mossambicus. PLOS ONE 12, e0177016 (2017).

22. Morimoto, H. et al. Procollagen c-proteinase enhancer-1 (pcpe-1) interacts with beta2-microglobulin (beta2-m) and may help initiate beta2-m amyloid fibril formation in connective tissues. Matrix Biol 27, 211–219 (2008).

23. Gibieža, P. & Prekeris, R. Rab gtpases and cell division. Small GTPases 9, 107–115 (2018).

24. Mi, H., Muruganujan, A., Casagrande, J. & Thomas, P. Large-scale gene function analysis with the panther classification system. Nat. Protoc. 8, 1551–1566 (2013).

25. Fernández-Fernández, M. R. & Valpuesta, J. M. Hsp70 chaperone: a master player in protein homeostasis. F1000Res 7 (2018).

26. Konstantinidis, G., Moustakas, A. & Stournaras, C. Regulation of myosin light chain function by BMP signaling controls actin cytoskeleton remodeling. Cell. Physiol. Biochem. 28, 1031–1044 (2011).

27. Cunningham, K. E. & Turner, J. R. Myosin light chain kinase: pulling the strings of epithelial tight junction function. Ann N Y Acad Sci 1258, 34–42 (2012).

28. Zhang, B. & Horvath, S. A general framework for weighted gene co-expression network analysis. Stat Appl Genet. Mol Biol 4, Article17 (2005).

29. Friedman, J., Hastie, T. & Tibshirani, R. Regularization paths for generalized linear models via coordinate descent. J. Stat. Softw. 33, 1–22 (2010).

30. Givens, G. & Hoeting, J. Computational statistics (John Wiley & Sons, New Jersey, 2013), 2 edn.

31. Gysi, D. M., Voigt, A., Fragoso, T. d. M., Almaas, E. & Nowick, K. wto: an r package for computing weighted topological overlap and a consensus network with integrated visualization tool. BMC Bioinforma. 19, 392 (2018).

32. Ravasz, E., Somera, A. L., Mongru, D. A., Oltvai, Z. N. & Barabasi, A. L. Hierarchical organization of modularity in metabolic networks. Science 297, 1551–1555 (2002).

33. Oldham, M. C. et al. Functional organization of the transcriptome in human brain. Nat Neurosci 11, 1271–1282 (2008).

34. Voigt, A. & Almaas, E. Assessment of weighted topological overlap (wto) to improve fidelity of gene co-expression networks. BMC BioInformatics 28, 58 (2019).

35. Benjamini, Y. & Hochberg, Y. Controlling the false discovery rate: a practical and powerful approach to multiple testing. J R Stat Soc Ser. B Stat Methodol 57, 289–300 (1995).

36. Derrida, B. & Flyvbjerg, H. Statistical Properties of Randomly Broken Objects and of Multivalley Structures in Disordered Systems. J Phys. 20 (1987).

